# Lipid Interactions of a Ciliary Membrane TRP Channel: Simulation and Structural Studies of Polycystin-2 (PC2)

**DOI:** 10.1101/589515

**Authors:** Qinrui Wang, George Hedger, Prafulla Aryal, Mariana Grieben, Chady Nasrallah, Agnese Baronina, Ashley C.W. Pike, Jiye Shi, Elisabeth P. Carpenter, Mark S.P. Sansom

## Abstract

Polycystin-2 (PC2) is a member of the TRPP subfamily of TRP channels and is present in ciliary membranes of the kidney. PC2 can be either homo-tetrameric, or heterotetrameric with PC1. PC2 shares a common transmembrane fold with other TRP channels, in addition to having a novel extracellular domain. Several TRP channels have been suggested to be regulated by lipids, including phosphatidylinositol phosphates (PIPs). We have combined molecular dynamics simulations with cryoelectron microscopy to explore possible lipid interactions sites on PC2. We propose that PC2 has a PIP-binding site close to the equivalent vanilloid/lipid-binding site in the TRPV1 channel. A 3.0 Å cryoelectron microscopy map reveals a binding site for cholesterol on PC2. Cholesterol interactions with the channel at this site are further characterized by MD simulations. These results help to position PC2 within an emerging model of the complex roles of lipids in the regulation and organization of ciliary membranes.

## Introduction

Ion channels are of considerable importance in numerous aspects of cell physiology, and mutations in channels cause many human diseases (Bagal et al., 2013). The transient receptor potential (TRP) superfamily of non-selective cation channels is a major class of ion channels found in all eukaryotes. They are involved in many aspects of cellular function, including thermosensation, osmotic pressure regulation, mechanosensation, and detection of noxious substances such as resiniferatoxin (Vetter and Lewis, 2011). TRP channels are activated and inhibited by a range of mechanisms, in response to thermal or chemical stimuli and/or mechanical forces (Venkatachalam and Montell, 2007). TRP channel mutations have been implicated in a number of different diseases (Nilius and Owsianik, 2010) and consequently are of interest as potential drug targets (Moran, 2018). Several TRP channels are modulated by membrane lipids (see for example (Fine et al., 2018; Wilkes et al., 2017; Yin et al., 2019), suggesting the possibility of lipid based therapies (Ciardo and Ferrer-Montiel, 2017).

Mammalian TRP channels may be divided into five families, the TRPC (Classical or Canonical), TRPV (Vanilloid), TRPM (Melastatin), TRPP (Polycystin), TRPML (Mucolipin) and TRPA (Ankyrin) subfamilies (Rohacs, 2009). Structurally, all TRP channels have a tetrameric architecture assembled from identical or similar subunits. Each of the four subunits is composed of six trans-membrane (TM) helices (S1-S6) with a pore loop region between S5 and S6 (Cao et al., 2013; Gao et al., 2016; Grieben et al., 2017; Huynh et al., 2016; Jin et al., 2017; Liao et al., 2013; Paulsen et al., 2015; Saotome et al., 2016; Shen et al., 2016; Su et al., 2018b; Wilkes et al., 2017; Zubcevic et al., 2016). TM helices S1 to S4 form a voltage sensor-like domain (VSLD), which is packed against the pore domain (S5-Pore-S6) of the adjacent chain, as is also seen in Kv channels, where structural data reveal that lipids mediate interactions of the voltage sensor domain with the pore (Long et al., 2007).

As a member of the TRPP subfamily, the polycystin-2 (PC2, also known as PKD2 or TRPP1) homo-tetramer has the same fold as other TRP channels. A novel extracellular domain (referred to as the TOP domain) is formed from a 218-residue insertion between S1 and S2, and a 20 residue insertion between S3 and S4 of the VSLD (Grieben et al., 2017; Shen et al., 2016; Wilkes et al., 2017). Structures of the closely related TRPP2L1 protein (Hulse et al., 2018; Su et al., 2018b), and also of the PC1/PC2 1:3 hetero-tetrameric complex (Su et al., 2018a) have also recently been determined.

Mutations in PC2 are responsible for about 15% of autosomal dominant polycystic kidney disease (ADPKD), which is one of the most prevalent genetic disorders in human, affecting 4 to 6 million people worldwide (Wilson, 2004). Most other cases of ADPKD (70%) are caused by mutations in PC1, which forms a 1:3 complex with PC2 by replacing one PC2 subunit in the channel region (Su et al., 2018a). Phenotypically, ADPKD is characterized by formation of fluid-filled renal cysts, which leads to progressive cystic enlargement in both kidneys and ultimately kidney failure (Pavel et al., 2016). Cysts or diverticula also frequently develop in intestines, liver and pancreas (Wilson, 2004). In addition, ADPKD is associated with increased risk of cardiovascular dysfunctions including aortic aneurysms, hypertension and heart-valve defects (Boucher and Sandford, 2004). Genetic data analysis has shown that there are at least 278 mutations in PC2 associated with ADPKD (see http://pkdb.pkdcure.org) (Gout et al., 2007). However, the underlying mechanisms by which these mutations lead to ADPKD are still poorly understood.

PC2 is widely distributed with relatively high expression in tubules within kidney cells (Chauvet et al., 2002), and was originally proposed to contribute to the transduction of extracellular mechanical stimuli caused by bending of the cilia into intracellular Ca^2+^ signals in the primary cilia of kidney epithelium (Nauli et al., 2003). In addition to its role in primary cilia, it has been proposed that PC2 could function as an intracellular Ca^2+^ release channel in the endoplasmic reticulum membrane (Koulen et al., 2002). More recent studies have suggested that Ca^2+^ signalling may not be involve in ciliary mechanosensation (Delling et al., 2016), and ciliary PC2 is a non-selective Na^+^, K^+^ channel, rather than a Ca^2+^ channel, as it is selective for Na^+^ and K^+^ over Ca^2+^ (Liu et al., 2018). PC2 and PC2-like channels have been identified across a wide range of organisms, from yeast to humans, highlighting the importance of this class of proteins (Shen et al., 2016). The membranes of primary cilia have a complex organization, including differences in membrane lipid composition between the base and the main body of the cilium (Garcia et al., 2018). Deletion of PC2 in mice leads to the formation of cilia that are 5-fold longer than normal cilia. Given the wider involvement of membrane lipid composition in regulation and signalling in primary cilia (Tsuji et al., 2019) and observations of lipid-like density in cryoelectron microscopy (cryoEM) structures of PC2 (Wilkes et al., 2017) it is therefore important to establish how PC2 may interact with specific lipids in its membrane environment.

Phospholipids have been extensively studied as modulators of membrane proteins. A number of lipids, and in particular phosphatidylinositol 4,5-bisphosphate (PIP_2_), have been shown to interact with and regulate ion channels (Basak et al., 2017; Hansen, 2015). G protein–coupled receptors (GPCRs) have been shown to be allosterically regulated by anionic lipids including phosphatidylserine (PS) and phosphatidylinositol (PI) (Dawaliby et al., 2016). Combined mass spectrometry, biochemical, and molecular dynamics simulation studies have revealed functionally important interactions of PIP_2_ with a number of Class A GPCRs (Yen et al., 2018). Perhaps the best studied example of PIP_2_ regulation of a membrane channel protein is provided by the inward rectifying potassium (Kir) channels, which are activated by PIP_2_ (Hansen, 2015; Shyng et al., 2000). PIP_2_ regulation has been suggested for almost all subfamilies of TRP channels, despite their possibly diverse activation mechanisms in response to different stimuli (Brauchi et al., 2007; Rohacs, 2007; Rohacs, 2009; Steinberg et al., 2014). Positive regulation by PIP_2_ has been indicated in at least five ion channels of the TRPM subfamily (Daniels et al., 2009; Liu and Liman, 2003; Runnels et al., 2002; Toth et al., 2015; Zhang et al., 2005). For TRPV channels, activation by PIP_2_ in excised patches has been reported for TRPV1 (Klein et al., 2008), TRPV5 (Lee et al., 2005) and TRPV6 (Thyagarajan et al., 2008). Indirect inhibition of TRPV1 by PIP_2_ in intact cells has also been reported (Thyagarajan et al., 2008). Recent cryoEM structures have revealed PIP_2_ binding sites on TRPV5 (Hughes et al., 2018b), TRPML1 (Fine et al., 2018), and TRPM8 (Yin et al., 2019). Given the importance of PIP_2_ distribution and dynamics in the membranes of primary cilia (Nakatsu, 2015), we decided to explore possible interactions of this anionic lipid with PC2. Cholesterol has also been shown to play a key role in signalling in ciliary membranes (Luchetti et al., 2016) and possibly in their organization (Garcia et al., 2018). Cholesterol is known to interact with many ion channels (Levitan et al., 2014; Morales-Lazaro and Rosenbaum, 2017) and receptors (Lee, 2018; Oates and Watts, 2011), so we decided to explore its possible interactions with PC2.

Molecular dynamics (MD) simulations provide an important ‘discovery tool’ for lipid interactions with membrane proteins (Corradi et al., 2018; Hedger and Sansom, 2016). For example, they have been used to predict PIP_2_ binding sites on K_ir_ channels (Schmidt et al., 2013; Stansfeld et al., 2009) and GPCRs (Yen et al., 2018). Given the growing number of structures of ion channels, binding affinities and specificity of interactions with lipids can be studied *in silico* via MD simulations (Domański et al., 2017; Hedger et al., 2019; Hedger et al., 2016)] to provide an indication of possible mechanisms of activation and allosteric modulation of channels by lipids.

Here we identify and characterise PIP_2_ interactions with PC2 by a combination of multiscale MD simulations with structural (cryoEM) and biochemical experiments. We propose there is a PIP_2_ binding site close to the equivalent vanilloid/lipid-binding site in TRPV1 channel (Gao et al., 2016). We also propose a binding site for cholesterol in PC2. These results thus help us to position PC2 within the emerging model of the complex roles of lipids in the regulation and organization of ciliary membranes (Weiss et al., 2019).

## Results & Discussion

### A phospholipid interaction site revealed by simulations

An initial exploration of possible lipid interaction sites on the transmembrane domain of PC2 was made using atomistic molecular dynamics simulations in which the PDB ID: 5K47 PC2 structure (as a representative of several PC2 structures, see following) was embedded in a lipid bilayer made up of a single species of phospholipid (palmitoyl-oleyl-phosphatidylcholine, PC; Fig. 1A). This process was repeated for three available molecular structures of wild-type PC2 (PDB ID: 5K47, 5MKF, 5T4D) and for a gain-of-function mutant (F604P) of PC2 (PDB ID: 6D1W) (Zheng et al., 2018b), yielding a total of more than 2 µs of atomistic simulations of PC2 in a PC bilayer. The simulations were examined in terms of regions of high probability density of occurrence of lipid molecules on the protein surface. In all 12 simulations (i.e. 3 repeats for each of the four structures, PDB ID: 5K47, 5MKF, 5T4D, 6D1W), high lipid occurrence densities (Fig. 1B) were observed in a pocket exposed to the intracellular leaflet of the lipid bilayer, between transmembrane helices S3, S4 and S5 (Fig. 1C), corresponding to one PC lipid molecule bound to each subunit of the PC2 tetramer. These results are illustrated for 5K47 in Fig. 1BC and similar results for 5MKF and 5T4D are shown in the Supporting Information, Fig. S1. The hydrophobic side chains from residues in S3, S4 and S5 create a hydrophobic pocket, within which the acyl tails of the bound lipid molecules reside. The phosphate oxygens of the bound lipid formed hydrogen bonds to the indole ring of Trp507 in S3 and to the hydroxyl group of Ser591 in the S4-S5 linker (SI Fig. S2). Thus, in a simple model lipid bilayer, we observe a phospholipid binding site on PC2 which is close to the proposed lipid and vanilloid (e.g. capsaicin) binding sites observed in cryoEM studies of the related TRPV1 and TRPV2 channels (Gao et al., 2016). Lipid or detergent binding has been observed at a similar location in cryo-EM structures of TRPV6 (McGoldrick et al., 2018; Singh et al., 2018), TRPV5 (Hughes et al., 2018b), TRPC3 (Fan et al., 2018), TRPC4 (Vinayagam et al., 2018), TRPM4 (Autzen et al., 2018; Duan et al., 2018b), TRPML3 (Hirschi et al., 2017) and TRPM7 (Duan et al., 2018a). This site has also been suggested to be the binding site for the TRPV5 inhibitor econazole (Hughes et al., 2018a), and for the TRPML1 lipid agonist PI(3,5)P_2_ (Chen et al., 2017).

**Figure 1:**
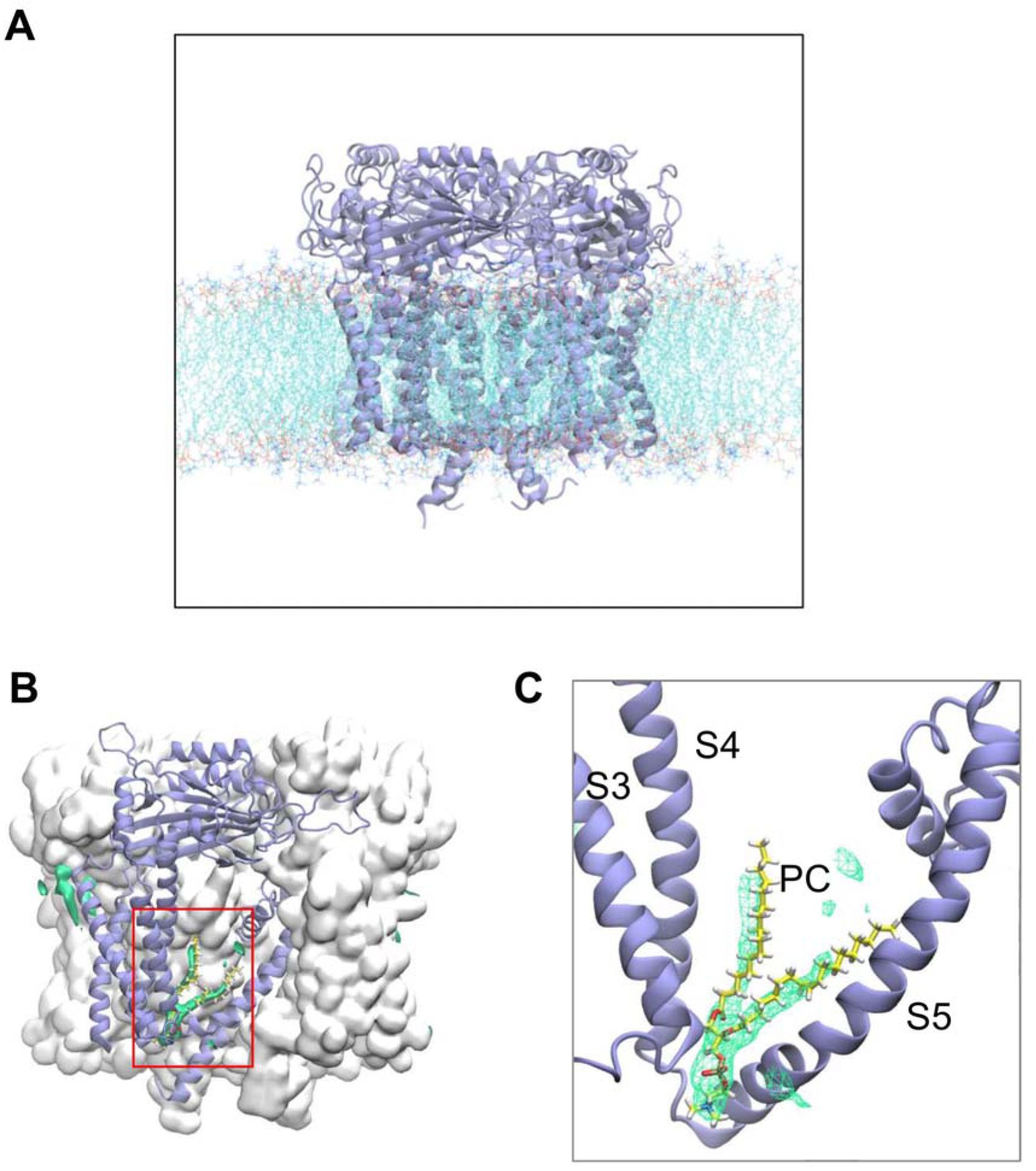
A possible phospholipid interaction site identified in atomistic simulations of PC2 in a PC bilayer. **A** The PC2 channel (PDB id 5K47) embedded in embedded in a phospholipid bilayer, shown using a snapshot from an atomistic simulation of the protein (colour) in a phosphatidylcholine (PC) bilayer. (Similar results for 5MKF and 5T4D are shown in the Supporting Information, Fig. S1). The lipid tails are in cyan, phosphates in orange and red, and choline nitrogens in blue. Water molecules are omitted for clarity. **B** The PC2 protein is shown as a grey surface, viewed perpendicular to the central pore axis, with one subunit depicted as a pale purple cartoon. Green isocontour surfaces represent a high probability of occurrence of phospholipid molecules. **C** Zoomed in view (red box in **B**) of the S3/S4/S5 pocket and the high phospholipid occurrence density with a PC molecule (taken from a simulation snapshot) shown within the density.

We subsequently used coarse grained (CG) simulations of PC2 in a lipid bilayer containing multiple lipid species (Ingolfsson et al., 2014; Koldsø et al., 2014) to explore the possible specificity of the lipid binding site. We embedded PC2 in an *in vivo*-mimetic bilayer (Fig. 2A), which contained the anionic lipids phosphatidylserine (PS) and PIP_2_ in the inner leaflet, a glycolipid (GM3) in the outer leaflet, and cholesterol (CHOL) in both leaflets of the bilayer. The lipid composition of the *in vivo*- mimetic bilayer membrane provided an approximation to a mammalian plasma membrane (Koldsø et al., 2014; Sampaio et al., 2011). Thus, the outer (i.e. extracellular or organelle luminal) leaflet contained PC:PE:SM:GM3:CHOL = 40:10:15:10:25; and the inner (i.e. intracellular) leaflet contained PC:PE:PS:PIP_2_:CHOL = 10:40:15:10:25). This provides an overall PIP_2_ concentration of 5% which is within the physiological range for a mammalian plasma membrane (Sampaio et al., 2011; van Meer et al., 2008).

**Figure 2:**
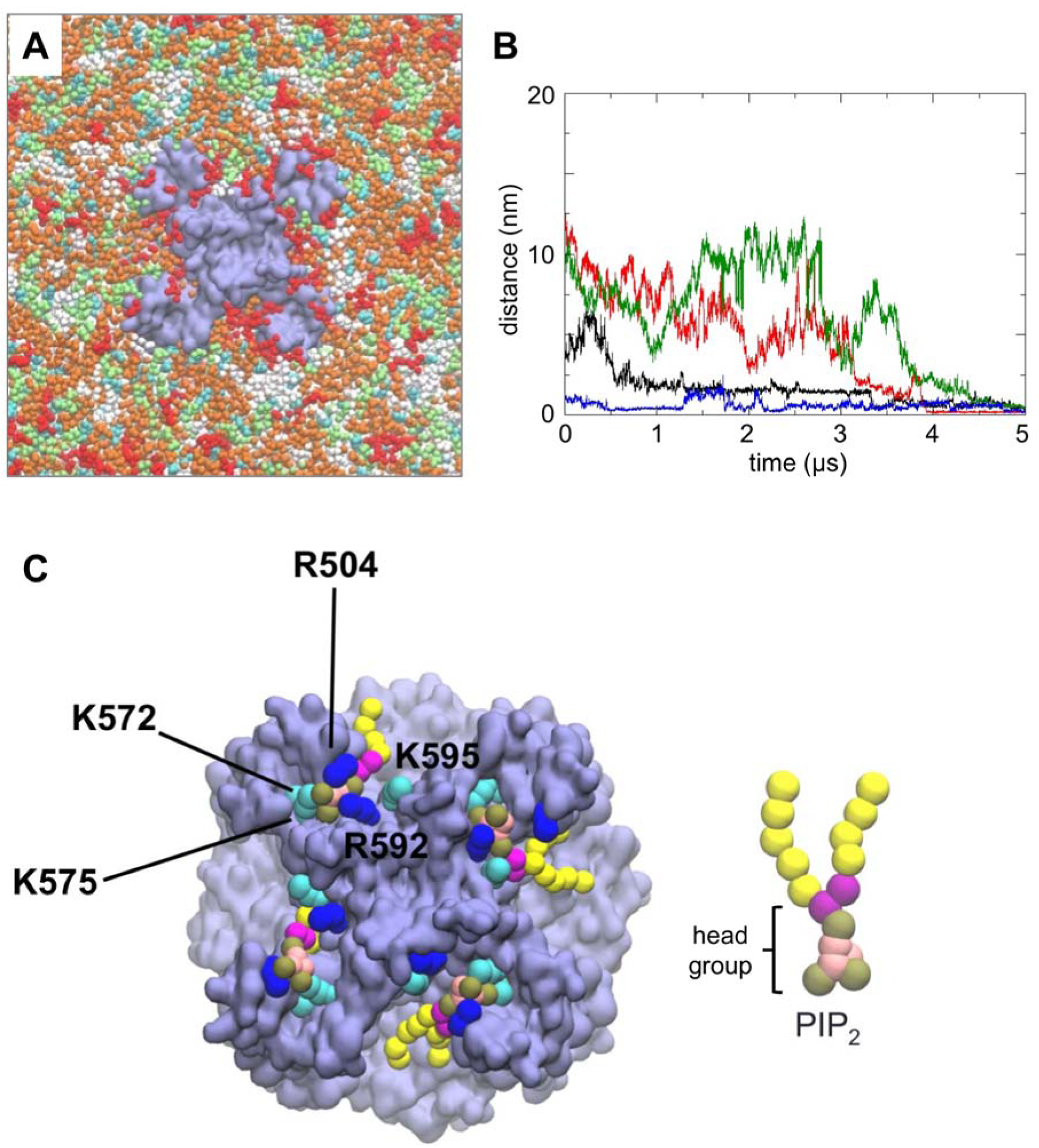
Coarse-grained (CG) simulations of PC2 in a *in vivo* mimetic mixed lipid bilayer. **A** PC2 (pale purple) in a mixed lipid bilayer, viewed from the intracellular face and showing molecules of PIP_2_ (red), cholesterol (cyan), PC (white), PE (orange) and PS (lime) in the inner leaflet of the bilayer. **B** Distance from the binding site as a function of time for four PIP_2_ molecules which bind to PC2 during a simulation in a mixed lipid bilayer. The distance is from the centre of mass of the headgroup of each PIP_2_ molecule to the centre of mass of the S505 and S591 sidechains at the site to which that lipid molecule eventually binds. The four colours correspond to the four different PIP_2_ molecules which eventually bind. **C** Snapshot from the end of the simulation shown in **A** showing PIP_2_ molecules (headgroup phosphate particles in brown, as shown in the CG representation of PIP_2_ on the right) at four sites on the channel tetramer. Sidechain particles of basic residues at the PIP_2_ binding site are shown in blue (arginine) and cyan (lysine).

Three independent simulations (each of 5 μs duration) of a single PC2 channel (structure 5K47) inserted in an *in vivo*-mimetic lipid bilayer with different random distributions of lipid molecules were performed. Simulations were also performed for the constitutively active F604P PC2 mutant (PDB ID: 6D1W). In each of the simulations, PIP_2_ molecules diffused in the bilayer (on a timescale of microseconds; Fig. 2B, SI Fig. S3, S4 & SI movie S1) resulting in random encounters with the channel molecule followed by binding to the previously identified sites on PC2, as demonstrated by tracking the distance *vs.* time of the PIP_2_ molecules from their eventual binding sites (Fig. 2B). From a final snapshot of one simulation (Fig. 2C), it can be seen that a PIP_2_ molecule has bound to each of the four sites on PC2. The head groups of the PIP_2_ molecules each interact with up to five basic residues in the S3/S4/S5 region: Arg504, Lys572, Lys575, Arg592 and Lys595, which are highly conserved between PC2 in different species (SI Fig. S5) This interaction is persistent in both wild type and mutant PC2. Given the presence of other negatively charged lipid species (PS) in the bilayer and of positively charged patches on the protein surface, the observation that PIP_2_ molecules were able to bind to the intracellular binding site suggests that this is specific for PIP_2_ and/or related lipids.

### Free energy landscapes for PC2/lipid interactions

To explore the energetics and selectivity of the PIP_2_/intracellular site interaction, we calculated potentials of mean force (PMFs) based on CG simulations. This approach has been used to explore interactions of anionic lipids (e.g. cardiolipin and PIP_2_) with a number of transporters (e.g. ANT1 (Hedger et al., 2016)), ion channels (e.g. K_ir_ channels (Domański et al., 2017)) and receptors (e.g. class A GPCRs (Song et al., 2019; Yen et al., 2018)). The PMF calculation provides a one dimensional free energy landscape for PIP_2_/PC2 interactions, and allows us to define the lipid specificity of this interaction (Fig. 3AB).

**Figure 3:**
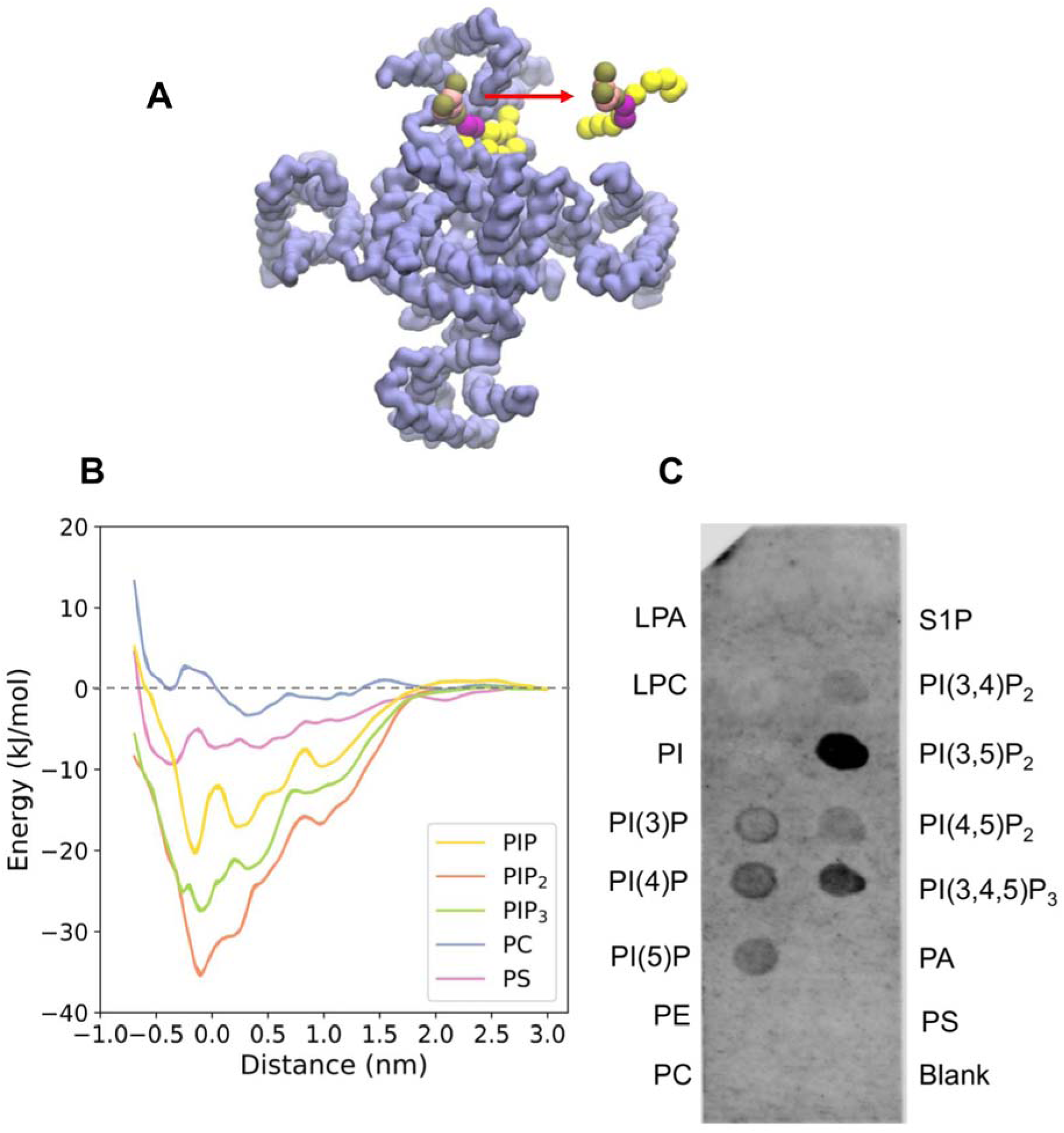
Free energy landscape for lipid interactions at the binding site on PC2. **A** Schematic representation of the reaction coordinate corresponding to the distance between centres of mass (COMs) of the headgroup of a lipid molecule (red) and of the two serines (S505, S591) within the lipid binding sites. **B** Potentials of mean force (PMFs) of the interactions of PIP_2_ (orange), PIP_3_ (green), PI(4)P (yellow), PS (purple) and PC (blue) with the lipid binding site on PC2. **C** PIP-strip data for interactions of different phospholipid species with PC2. Note the null results for the anionic PA, PS, and S1P lipids, and for the zwitterionic PE and PC controls.

To estimate the free energy landscapes for phosphatidylinositol monophosphate (PIP), PIP_2,_ phosphatidylinositol trisphosphate (PIP_3_), PC, and of PS (Fig. 3B) interacting with PC2 at the intracellular site defined by S3/S4/S5, CG models of a PC2 channel molecule embedded in a PC bilayer with a PIP, PIP_2_ or PIP_3_ molecule inserted into each of the four binding sites were used. For computational efficiency, the PC2 structure was truncated, removing the TOP domain (Ser244 – Leu462). The initial configurations were lipid bound states. One of the bound lipid molecules was pulled away from the binding site to assess the free energy of interaction. A one-dimensional reaction coordinate was defined as the distance in the plane of the bilayer between the centres of mass of the lipid headgroup and two serines (Ser505, Ser591) within the lipid binding site of PC2 (Fig. 3A). The free energy profile (i.e. PMF) for PIP_2_ shows a clear minimum close to the initial position of the PIP_2_ molecule in the binding site, with a well depth of −37 kJ/mol (Fig. 3B). Comparison of PMF profiles as a function of window simulation duration suggested that convergence had been achieved within 1.5 µs per window; see SI Fig. S6). PIP and PIP_3_ exhibit weaker binding to PC2 with free energies of −28 kJ/mol and −20 kJ/mol respectively. In contrast, the anionic lipid PS showed significantly weaker binding to PC2 with a well depth of approximately −9 kJ/mol, indicating a clear selectivity of the binding site for PIP species over PS. Phosphatidyl choline (PC) shows even weaker interactions, as would be anticipated given the PMF is evaluated for a single lipid molecule in an environment of a PC bilayer. The two minima in PMF for the PC are likely to reflect annular shells of relatively immobilized lipid molecules around the PC2 channel (see SI Fig. S7), as have been seen for a number of channel and other membrane proteins in simulations (Goose and Sansom, 2013; Niemela et al., 2010).

The predicted binding of PIP_2_ to PC2 was tested biochemically using PIP-strips (Shirey et al., 2017), nitrocellulose membranes with lipids including phosphoinositides and other anionic and zwitterionic phospholipids spotted onto their surface (Eshelon Biosciences Ltd). Protein binding to relevant lipids can be detected with antibodies to the tagged protein. This method has been used for example to confirm binding of PIP_2_ to the GluA1 ionotropic glutamate receptor (Seebohm et al., 2014). The results from this assay (*N* = 5 biological repeats; see SI Fig. S8) suggest that detergent-solubilized truncated PC2 can bind a range of phosphatidylinositol phosphates (PIPs), including PI(4,5)P_2_. In contrast, it does not exhibit interactions with simple anionic (PS, PA, SP1) or zwitterionic (PE, PC) lipids under the experimental condition used. It has been shown that the basal regions of primary cilia membranes contain PIP_2_ whilst the upper regions of the ciliary membrane have an elevated level of PI(4)P (Garcia et al., 2018; Nakatsu, 2015). Thus, it is likely that PC2 binds to PIP_2_ *in vivo*. In this context it is of interest that OCLR1, a lipid phosphatase that converts PI(4,5)P_2_ to PI(4)P, modulates the length of cilia in renal epithelial cells and loss of its function in Lowe syndrome is associated with progressive renal malfunction (Rbaibi et al., 2012).

### The PIP_2_ binding site

Having established that PC2 binds PIP_2_ selectively, we examined the interactions of the protein with the lipid in more detail, based on a CG snapshot structure of the PC2-PIP_2_ complex corresponding to the energy minimum in the PMF. As noted above, the head group of PIP_2_ interacts with five basic residues: Arg504, Lys572, Lys575, Arg592 and Lys595. In particular, the 1 ′ phosphate interacts closely with Arg592, and the 4′and 5′phosphates interact with Lys572 and Lys575 respectively. A binding site for PIP_2_ formed by a cluster of basic residues is seen in other ion channels, e.g. Kir channels (Hansen et al., 2011; Lacin et al., 2017), and in GPCRs (Song et al., 2019; Yen et al., 2018).

To evaluate the PIP_2_ binding site in more detail, the CG structure corresponding to the final snapshot of the sampling window for the energy minimum configuration was converted into an atomistic representation. A two-stage atomistic simulation was then performed. Firstly, a short (30 ns) simulation was performed in which harmonic restraints were applied to the distances between the 1′, 4′, and 5′ phosphates of PIP_2_ and the sidechains of Arg592, Lys575 and Lys572 respectively, in order to relax the atomistic model whilst maintaining the interactions seen in the CG PMF calculations. The distance restraints were then removed and three replicates of an unrestrained simulation (durations 200 to 250 ns) were run to allow the PIP_2_ to explore the binding site on PC2. Final snapshots from the restrained and unrestrained simulations are shown (Fig. 4A). The interactions between the lipid head group and the interacting residues are seen to be dynamic and to vary stochastically between replicate simulations. Interactions of the tails are very dynamic. Thus, for residues Arg504, Lys575, Arg592 and Lys595, fluctuating numbers of hydrogen bonds were formed with PIP_2_ throughout the simulations (Fig. 4B and SI movie S2). Such fluctuations in PIP_2_ binding are not unique to PC2, they have also been reported for PIP_2_ molecules bound to K_ir_ channels, e.g. (Lacin et al., 2017).

**Figure 4:**
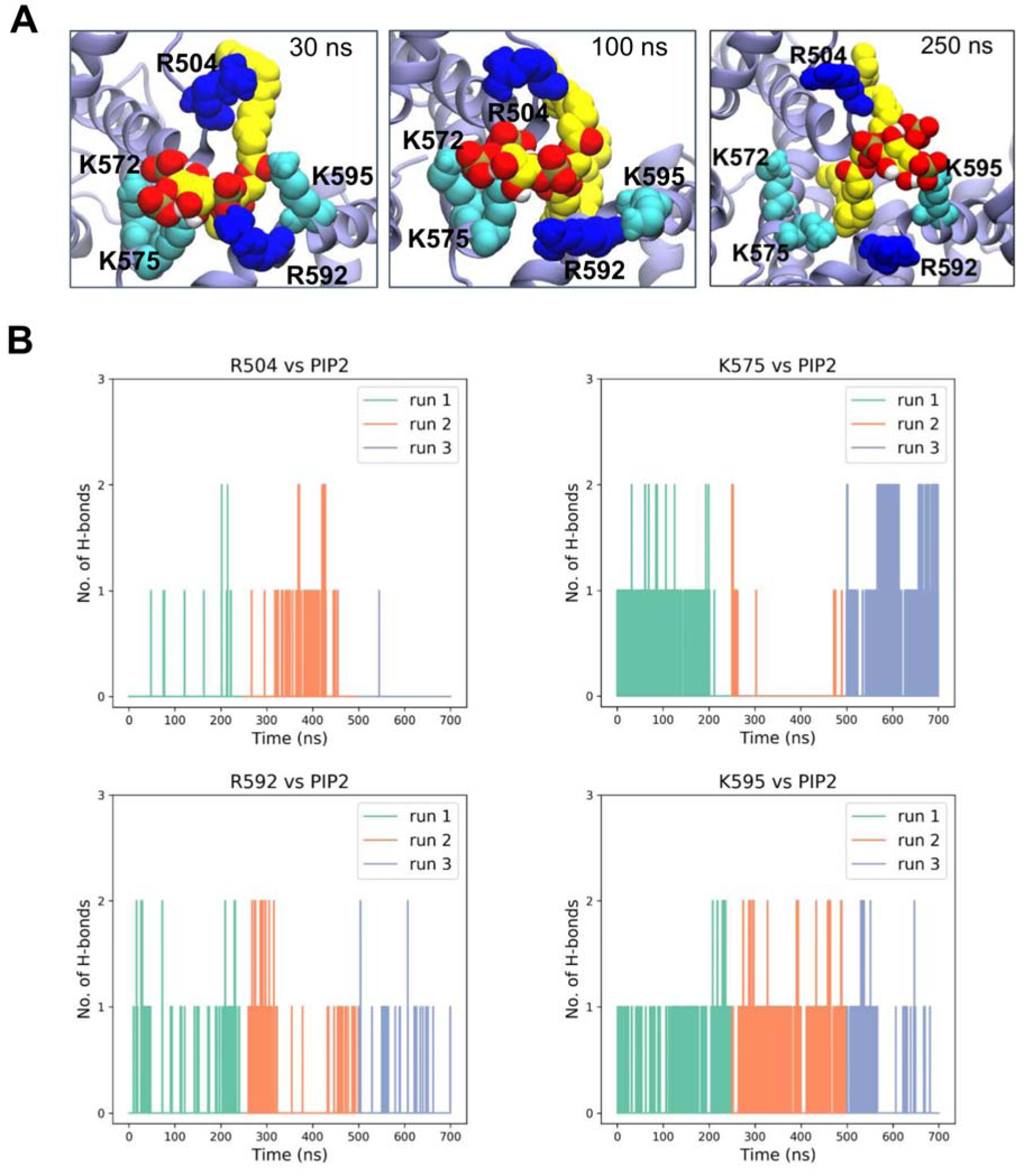
Protein-lipid interactions showing the head group of PIP_2_ coordinated by five basic residues (R504, K572, K575, R592 and K595) at the binding site. **A** Snapshot structure corresponding to the CG energy minimum was converted into an atomistic representation and used as the basis of 30 ns distance restrained simulations (see text for details) followed by 250 ns of unrestrained simulations (with snapshots at 100 and 250 ns). **C** H-bonding interactions between key basic sidechains and the PIP_2_ headgroup during 3 replicates (run 1 0-250 ns, run 2 250-500 ns; run 3 500-710 ns) of the unrestrained atomistic simulations.

### Cryoelectron microscopy

In order to investigate binding of PIP_2_ to PC2, we determined two cryoEM structures of PC2, in the presence of PI(4,5)P_2_ to 3 Å resolution and of PI(3,5)P_2_ to 3.4 Å. We used similar conditions to our original 4.2 Å PC2 structure (PDB ID: 5K47), with a truncated construct (residues Pro185 to Asp723), purified in the detergent UDM (n-undecyl-β-D-maltopyranoside). The overall conformation of the structures remained the same as our original structure, i.e. a closed conformation, although we did in this case observe a disulphide bond (between residues in the TOP domain) which was absent in our original structure. We inspected the region of the maps where PIP_2_ was found in the simulations, to see whether there is any evidence for density corresponding to lipids at this site. As can be seen (Fig. 5) there are non-protein densities located between S3, S4 and S5 in both structures, suggestive of lipid and/or detergent occupancy (Fig. 5C and SI Fig. S8). The density observed in the PI(3,5)P_2_ structure may correspond to a lipid molecule (Figure 6B), although the presumed headgroup region of the density is not clear enough to unequivocally interpret it as a PIP molecule. With the PI(4,5)P_2_ higher resolution structure, the density resembles two detergent molecules, consistent with the observation from the PIP strips that the PI(4,5)P_2_ appears to bind less well to PC2 and may not be able to displace high concentrations of detergent. With PI(3,5)P_2_ the density suggests a lipid molecule bound at this site, but there are few interactions between the region where the headgroup lies and the potential lipid and overall the lipid is insufficiently clear to unequivocally assign a lipid molecule to this density, so no lipid is placed at this site in the final PDB file. In TRP channel structures with lipid molecules built into similar locations to the site we observe (e.g. (Gao et al., 2016; Hughes et al., 2018b), the lipid headgroups are coordinated not only by residues on S3, S4 and S5 but also by residues in the cytosolic pre-S1 or post-S6 domains. The lack of cytosolic domains of our structures may contribute to the flexibility of the headgroup of the bound lipid, helping to explain why the densities in the constructs used for these structures are not well resolved in the headgroup region. Similarly, there is some undefined density in this region in structures obtained using soy extract polar lipids (which are likely to contain some PIs) (Shen et al., 2016). We inspected the cryoEM density map of the latter structure (https://www.emdataresource.org/EMD-8354), which was determined in nanodiscs (Shen et al., 2016). In the region of interest, there is density which could represent a bound phospholipid molecule. Together with our data, this suggests that the hydrophobic pocket identified in our simulations is a phospholipid binding site. In addition, with the more soluble and potentially higher-affinity version of PIP_2_, there is a small outward movement (∼ 2 Å) of the VSLD at the S2-S3 linker region and an inward movement (∼ 2 Å) towards the central axis of the cytosolic extension of S6 helix in the PI(3,5)P_2_ structure compared to the PI(4,5)P_2_ structure (which may correlate with the apparently stronger binding of PI(3,5)P_2_ seen in the PIP-strip assay). A distinct PIP_2_ site in TRP channels has been proposed, formed between the Pre-S1 and TRP-like domains (Zheng et al., 2018a) although the structural basis for this remains uncertain. However, a comparable PIP_2_ site was observed in TRPV5 (Hughes et al., 2018b), and a PIP_2_ site between helices S1, S2 and S3 was seen for TRPML1 (Fine et al., 2018). It seems possible that more than one PIP lipid binding site may be present on TRP channels (see also (Yin et al., 2019)), which could explain some of the apparent diversity of effects (both positive and negative) of PIPs on these channels (Rohacs, 2015).

**Figure 5:**
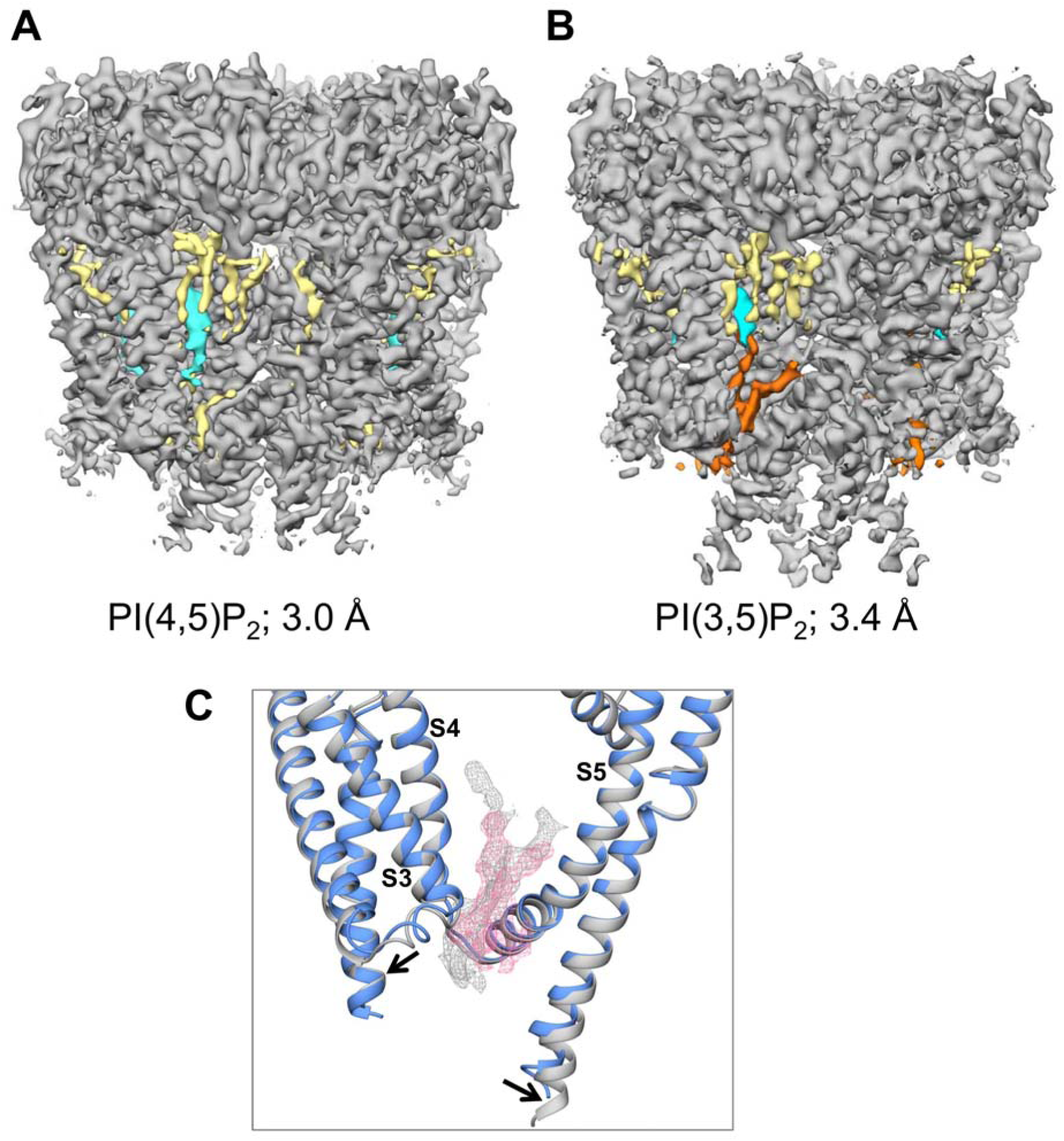
Lipid binding sites suggested by cryoEM maps of PC2 obtained **A** in the presence of PI(4,5)P_2_ (3.0 Å resolution) and **B** in the presence of PI(3,5)P_2_ (3.4 Å resolution). Protein density (contoured at 3.2 sigma) is in grey, detergent density in yellow, lipid density in orange, and cholesterol density in cyan. **C** Expanded view around the proposed lipid binding site between S3, S4 and S5 showing density from the 3.0 Å map (pink; see SI Fig. S9) and from the 3.4 Å map (grey).

**Figure 6:**
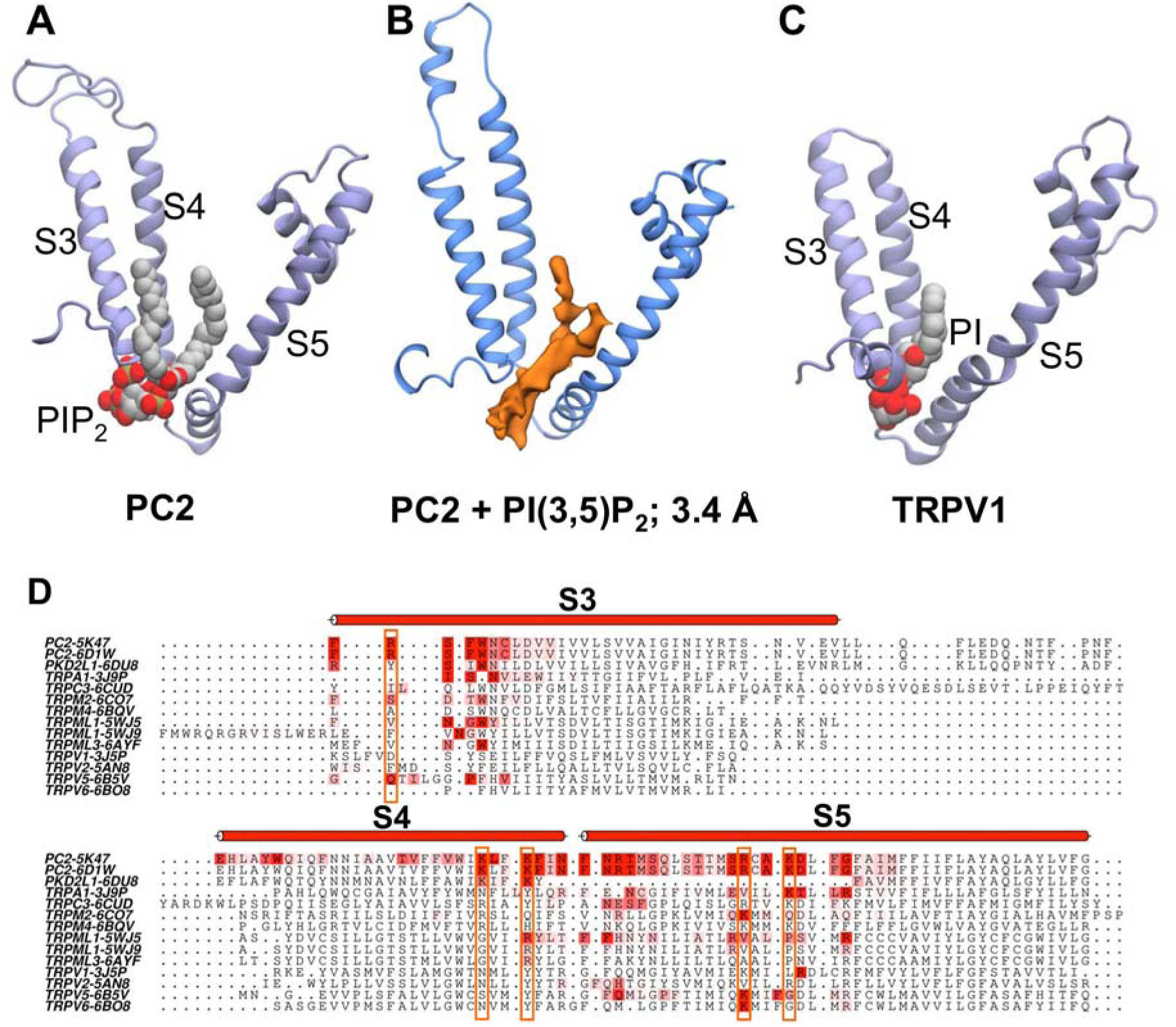
Comparison of PI lipids bound to PC2 and to TRPV1 with cryoEM density. **A** PIP_2_ bound to PC2 (as revealed by the current simulation study); **B** Lipid-like density in the cryoEM maps of PC2 obtained in the presence of PI(3,5)P_2_ (3.4 Å resolution; see Fig. 5); **C** PI bound to TRPV1 (as revealed by cryoelectron microscopy, PDB ID 5IRZ). In each case the lipid molecule or density is located between the S3, S4 and S5 helices of the VSLD. **D** A sequence alignment coloured on contacts with PIP_2_ in the mixed lipid simulations. Residues of the region around the binding pocket between the S3, S4 and S5 helices are coloured (on a white to red scale) according to the frequency of the interactions of PIP_2_ headgroups with each residue. The five basic residues of PC2 which form interactions with the headgroup of PIP_2_ at the binding site (i.e. R504, K572, K575, R592 and K595) are boxed.

### PIP interactions in TRP channels

The proposed PIP interaction site on the TM domain of PC2 may be compared with a lipid binding site, close to the vanilloid ligand-binding site, which has been seen in the structure of TRPV1 and which has been interpreted as being occupied by a phosphatidylinositol molecule (Gao et al., 2016). Comparison of the two sites (Fig. 6) reveals strong similarities, especially the location of the anionic phosphate-containing lipid headgroup at the N-terminal region of the S3 helix dipole (Hol et al., 1978). Furthermore, the predicted interaction site on PC2 and the PI site on TRPV1 both agree well with the density tentatively interpreted as a lipid molecule in our 3.4 Å cryoEM map (see above). To explore this possible common binding site further we extended our CG simulations of PC2 in a mixed lipid *in vivo*-mimetic to a number of different TRP channel structures (see Table 1). The results suggest (Fig. 6D) a degree of conservation of the proposed PIP-channel interaction between different families of TRP channels.

**Table 1:**
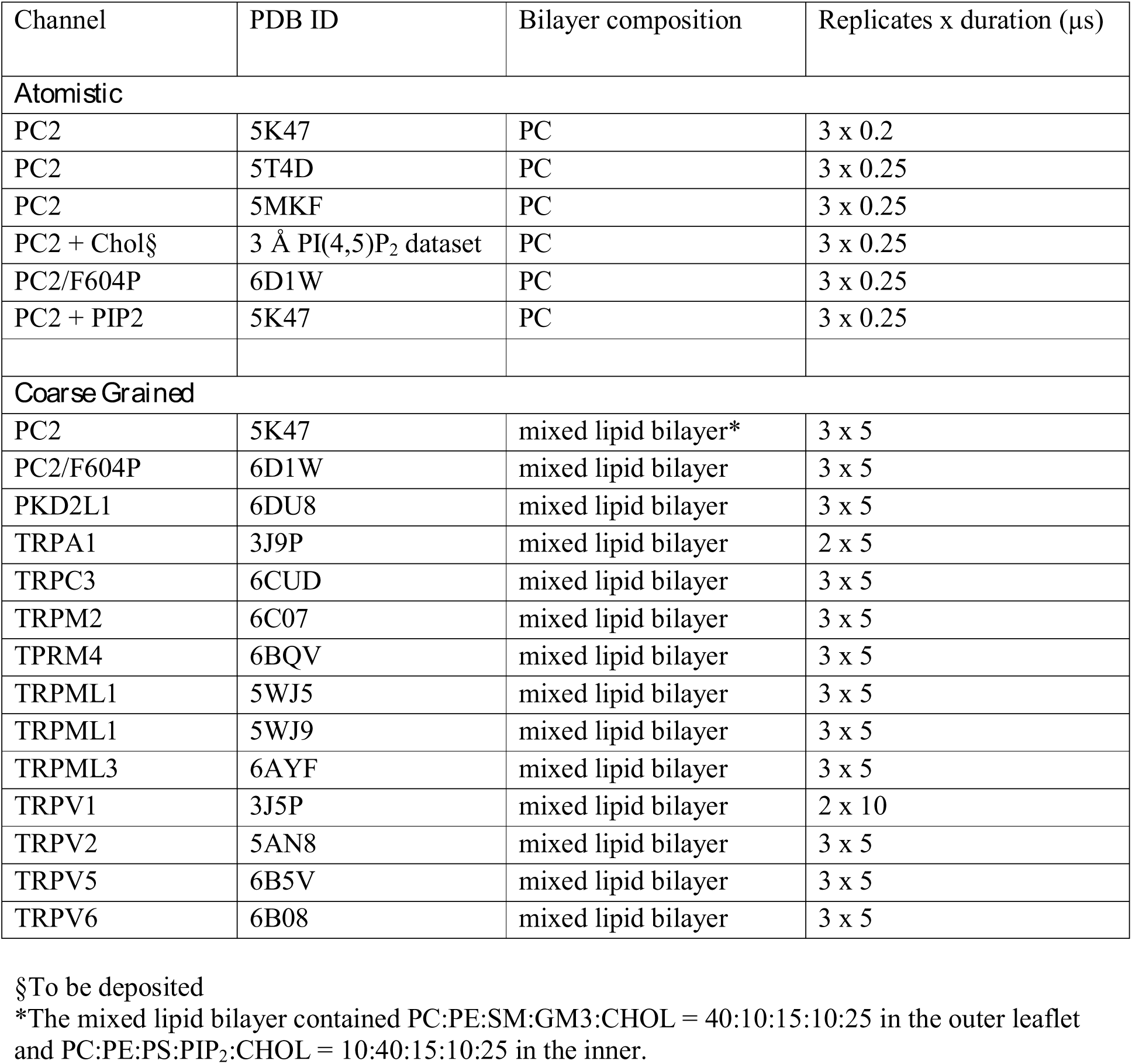
Summary of Simulations

### Cholesterol interactions

Examination of the cryoEM density also revealed a possible site for (co-purified) cholesterol (Fig. 7A) located between the S3 andS4 helices and S6 of the adjacent subunit. The cholesterol must have remained bound to PC2 throughout the extraction and purification process, since no cholesterol, or cholesterol mimetic such as cholesteryl hemi-succinate (CHS), was used in PC2 purifications. Interestingly comparable density may be observed for the 5T4D structure. Given the importance of cholesterol in the ciliary membranes (Garcia et al., 2018) we analysed our CG-MD simulations of PC2 in an *in vivo*-mimetic (i.e. mixed lipid) bilayer to identify possible interactions of cholesterol with the channel (Fig. 7B and SI Fig. S10). The results of these simulations revealed a cholesterol interaction site close to the site suggested by the cryoEM density. This indicates that this site may interact with cholesterol present in the bilayer. We note that recent comparable CG-MD simulations of GPCRs have identified interaction of cholesterol at sites which bind cholesterol in a number of crystal structures (Song et al., 2019). Having identified a cholesterol binding site, we performed CG PMF calculations to estimate the free energy landscape for the cholesterol interaction (Fig. 7D). This showed a weaker interaction than for PIPs (above), with a free energy well depth comparable to that for binding of cholesterol to other membrane proteins (see e.g. (Hedger et al., 2019)).

**Figure 7:**
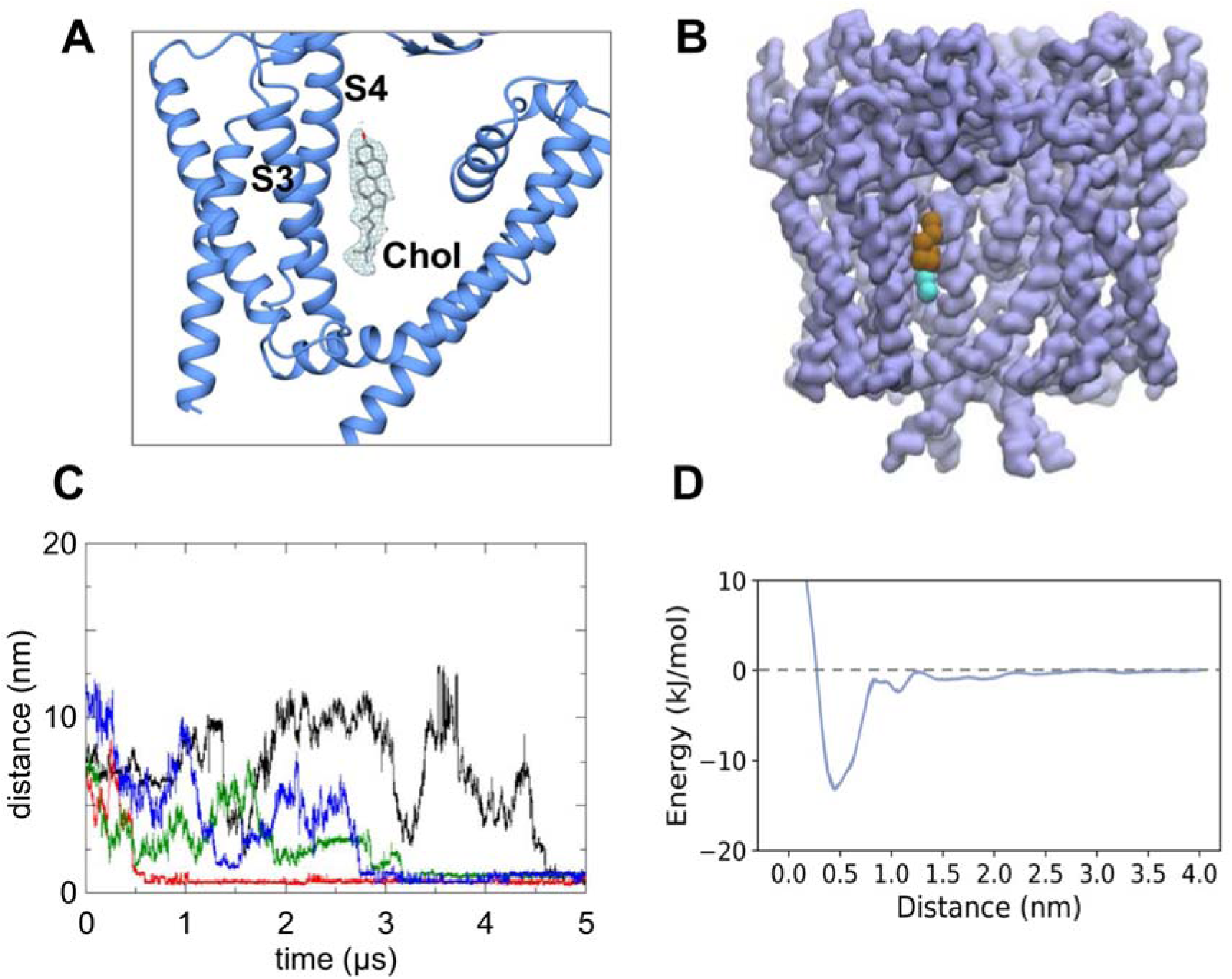
Cholesterol interactions with PC2. **A** CryoEM density (from the 3.0 Å map, contoured at 2.2 sigma; see Fig. 5A) corresponding to a binding site for cholesterol located between the S3 and S4 helices and helix S6 of the adjacent subunit. **B** Cholesterol observed to bind to the same site in CG simulations of PC2 in a *in vivo* mimetic mixed lipid bilayer (see Fig. 2). **C** Distance from the binding site as a function of time for four cholesterol molecules which bind to PC2 during a simulation in a mixed lipid bilayer. **D** Potential of mean force (PMF) for the interaction of cholesterol with the binding site on PC2.

To further test the possible cholesterol binding site, we performed atomistic simulations (3 × 250 ns) starting with a model of cholesterol built into the cryoEM density. From the simulations it is evident that cholesterol interacts dynamically at this site, which is located at the interface between the VSLD and S6 with the long axis of cholesterol running approximately parallel to helices S3 and S4. Thus, for the four symmetry-related binding sites on the channel, one site exhibits a stably bound cholesterol, one site has a cholesterol which transiently dissociates then rebinds, and the two other sites show intermediate behaviour (Fig. 8A & SI Fig. S11). This is consistent with a number of simulation studies of cholesterol on GPCRs which indicate relatively dynamic, loose binding (Fig. 8B; see (Hedger et al., 2019) for a more detailed discussion). The hydroxyl group mainly hydrogen bonded to Gln557 and Asn560. The steroid nucleus of cholesterol sits within a shallow hydrophobic pocket formed by a group of isoleucine (Ile561, Ile659), leucine (Leu517, Leu656) and valine (Val564, Val655) residues (Fig. 8C). This is consistent with previous studies of cholesterol binding in GPCRs, which showed that isoleucine is the main contacting residue within the binding sites (Gimpl, 2016; Hedger et al., 2019). The hydrocarbon chain of cholesterol showed considerable mobility throughout the simulation, which may explain the lack of density around this region in our cryoEM map. It is tempting to speculate about the possible biological importance of cholesterol interactions with PC2. For example, cholesterol plays a key role in signalling via the ciliary GPCR Smoothened, and it is possible that different regions of cilia differ in the cholesterol content of their membranes (Luchetti et al., 2016). However, further biochemical studies will be needed to establish a possible role of cholesterol in regulation of PC2 in ciliary membranes.

**Figure 8:**
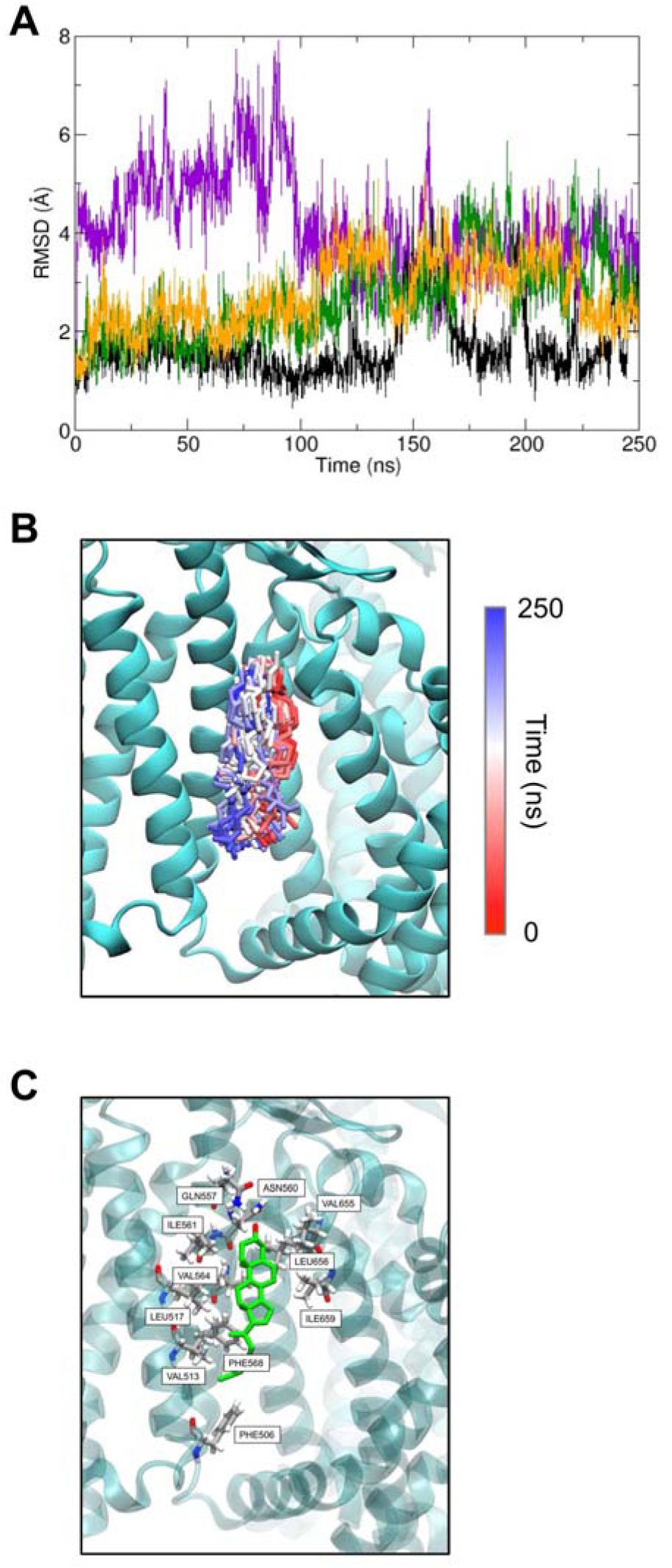
Atomistic simulations of cholesterol-bound PC2, with the cholesterol molecule initially built into the cryoEM density. **A** RMSD vs. time for the four cholesterol molecules bound to PC2 during a 250 ns atomistic MD simulation. The large fluctuations in RMSD for two molecules (purple and green curves) demonstrate the relatively loose binding of cholesterol at this site. **B** Snapshot structures of cholesterol (stick representation; taken every 1 ns) at the binding site on PC2. Each of these structures is coloured according to the corresponding simulation time on the RWB colour-scale shown, thus structures at the start of the simulation are coloured red, and those at the end in blue. **C** Final simulation snapshot (t = 250 ns) showing the arrangement of key binding site residues.

### Possible limitations

The current resolution achievable in many cryoEM studies of membrane proteins means that it is generally difficult to unambiguously identify bound lipids. Molecular simulations thus provide a valuable tool in assessing possible identities of bound lipids. In our studies, MD simulations are strongly suggestive of PIP_2_ and cholesterol binding at the two sites identified. However, the experimental data only allowed unambiguous confirmation of these predictions in the case of cholesterol, not PIP_2_. PIP strip assays were used to identify possible lipid species interacting with PC2, but quantification of relative binding affinities for different lipids could not be established. Similarly, in our PMF calculations, coarse graining of the lipid models does not allow distinction between different PIP species with the same charge. In addition, the PC2 construct used for PIP strips is truncated (as is the construct in the structural studies and the simulations). It is therefore uncertain whether full-length PC2 protein would exhibit the same lipid binding behaviour, especially for PIP where the binding site is close to the termini of the truncation.

## Conclusions

Using multiscale MD simulations, we identified a lipid binding pocket in the human TRP channel PC2 between the S3 and S4 TM helices and the S4-S5 linker. Free energy calculations suggested that this binding site is selective for PIP_2_ and related PIPs, and this is supported by biochemical, (biophysical), and structural data. The location of this site is analogous to the shared binding site for vanilloid ligands and phosphatidylinositol lipids in the canonical TRPV1 channel (Fig. 5AB). This region is distinct from the location between the VSLD and the core pore-forming domain) of the 5MKE structure of PC2 (Wilkes et al., 2017) where lipid-like density is modelled as phosphatidic acid and as palmitic acids binding within the extracellular leaflet region of the channel protein.

Structural and simulation studies have shown that the movement of the S4-S5 linker is important for gating in voltage-gated cation channels. For example, in the voltage-gated potassium channel Kv1.2, the constriction or dilation of the pore domain is controlled by the voltage sensor domain through the S4-S5 linker (Long et al., 2005). The transmembrane helices of TRP channels and voltage-gated cation channels share high structural similarity, indicating a possible common functional role of the S4-S5 linker. Indeed, it has been shown that TRPV1 (Gao et al., 2016; Yang et al., 2015), TRPV2 (Yang et al., 2016) and TRPV4 (Teng et al., 2015) use the same mechanism to gate the channels, and disruption of the interaction configurations at the linker region by ligands/lipids could change the gating probability.

A significant difference between the TRPV channels and the PC2 structures is that the binding sites in TRPVs are enclosed by the cytoplasmic C-terminal TRP domains, which is conserved in most TRP subfamilies. However, the TRP domain is not present in the TRPP subfamily and there are more than 200 residues missing from both the N- and C-termini in the PC2 structures. The PIP lipid head groups might be additionally coordinated in the presence of the intracellular domains. Sequence alignment shows that the basic residues involved in PIP_2_ binding (Lys572, Lys575, Arg592 and Lys595) are highly conserved (Fig. S5), which suggests that these residues may play a crucial role in modulating the channel activity. Disruption of the native intramolecular interactions involving these residues may also alter channel activity. Residue Asp511 is central to the hydrogen-bonding network involving Lys572 and Lys575. The naturally occurring mutation D511V has been well characterised to be channel inactivating (Koulen et al., 2002). The loss of electrostatic interactions between the Asp and Lys residues could allow Lys572 and Lys575 to have stronger interactions with PIP_2_, thus magnifying the inhibitory effect of PIP_2_ on PC2. The similar binding energy of PIP_2_ in PC2 and Kir2,2 (Domanski et al., 2017) channels is also supportive of the existence of a PIP_2_ binding site in PC2.

Our structural and simulation studies also suggest a cholesterol-binding site on the TOP domain, outer-leaflet facing surface of the PC2 molecule. This seems to be close to the site suggested to interact with a phospholipid (modelled as phosphatidic acid; see PDB: 2MKF and 2MKE) in the study of Wilkes et al. (Wilkes et al., 2017). It is distinct from but adjacent to the sites interpreted as corresponding to CHS in the latter study. Possible interactions of TRP channels with cholesterol have not been studied in detail, although some physiological data are available (Morales-Lazaro and Rosenbaum, 2017). Given the emerging importance of cholesterol interactions with other ion channels and with receptors (Bukiya and Rosenhouse-Dantsker, 2017; De Jesus-Perez et al., 2018; Lee, 2018) it would not be surprising if cholesterol interactions with TRP channels were of functional importance, although detailed studies of a number of TRP channel family members will be required to establish this. Establishing the interactions of TRP channels with lipids provides an avenue towards defining novel druggable sites on this important class of channel molecules.

## Methods

### Molecular Model Preparation

N-acetyl-D-glucosamines were removed from all protein structures, and missing side chains and loops were modelled using MODELLER version 9v10 (Fiser and Sali, 2003). Models were visually inspected. Structurally realistic models with the lowest value of the MODELLER objective function were chosen for subsequent simulations.

### CG Simulations

CGMD simulations were performed using GROMACS version 4.6 (www.gromacs.org) (Pronk et al., 2013) with the MARTINI version 2.1 force field (Monticelli et al., 2008). CG simulations within PMF calculations were done with GROMACS version 5.1 (www.gromacs.org) (Abraham et al., 2015) with the MARTINI version 2.2 force field (de Jong et al., 2013). An elastic network (Periole et al., 2009) was applied with elastic bond force constant of 500 kJ/mol/nm^2^ and an upper elastic bond cut-off of 0.9 nm. The standard MARTINI water beads with van der Waals radii of 0.21 nm were used to solvate all systems, which were then neutralised with NaCl at a concentration of 0.15 M. CG lipids and ions were described by MARTINI version 2.0 lipids and ions respectively.

Each simulation system contains only one copy of a protein. Initially, PC lipids were randomly placed around the proteins. Correct positioning of protein in a lipid bilayer was achieved by a 100 ns self-assembly simulation (Scott et al., 2008). The CG system for PDB ID: 5K47 was equilibrated for 400 ns after the initial self-assembly. Simulation systems for the other PC2 structures (i.e. PDB ID: 5MKF and 5T4D) were set up by aligning and replacing the protein in the PDB 5K47 POPC-only system, and then equilibrating for 1 μs. Systems used for mixed-lipid simulations were set up by exchanging PC molecules for other lipids using a locally developed script.

All CGMD simulations were performed at a temperature of 310 K and a pressure of 1 bar. V-rescale thermostat (Bussi et al., 2007) was used to maintain the temperature using a coupling time constant of τ_t_ = 1 ps. Protein, lipids, and solvent (water + ions) were coupled separately to the temperature bath. For self-assembly and equilibration simulations, pressure was controlled by semi-isotropic pressure coupling with the Berendsen barostat (Berendsen et al., 1984) with a compressibility of 5 × 10^-6^/bar and a time constant of τ_p_ = 4 ps. Production runs used a Parrinello-Rahman barostat with coupling constant of 12 ps and compressibility of 3 × 10^-4^/bar for pressure control.

### Atomistic Simulations

These were setup and run using a two-step multiscale procedure (Stansfeld et al., 2015; Stansfeld and Sansom, 2011), starting with a CG simulation to stably insert the protein in a lipid bilayer (see above), followed by conversion to an atomistic representation and subsequent atomistic simulations. A fragment based protocol (Stansfeld and Sansom, 2011) was used for the CG to atomistic conversion. For the PDB 5K47 system in a PC-only bilayer, the final frame of the 400 ns equilibration was converted to atomistic representation. Three repeats of 200 ns atomistic simulations with different initial random seeds were performed after a 6 ns unrestrained equilibration. Simulations were carried out using GROMACS version 4.6 with the CHARMM36 force field (Best et al., 2012). The water model used was TIP3P (Jorgensen et al., 1983). Atomistic systems for PDBs 5MKF and 5T4D were converted from the final frame of the 1 μs CG equilibration runs. Three repeats of 250 ns atomistic simulations were done for both structures after 1ns fully restrained equilibration.

Simulation systems of PC2 structures with bound cholesterol were setup using the CHARMM-GUI (www.charmm-gui.org). The protein with bound cholesterols were embedded in POPC-only bilayer. Simulations were performed using GROMACS version 5.1 with the CHARMM36m (Huang et al., 2017) force field and TIP3P water model (Jorgensen et al., 1983). Equilibration of the system was done in six steps with gradually decreasing restraint force constants on the protein and cholesterol molecules. Three repeats of 250 ns unrestrained atomistic simulations were then performed.

For all atomistic MD simulations, the long-range electrostatics (< 1 nm) was modelled with the Particle Mesh Ewald (PME) method (Essmann et al., 1995). Temperature coupling was done with V-scale thermostat at 310 K. The Parrinello-Rahman barostat (Parrinello and Rahman, 1981) with a reference pressure of 1 bat and a compressibility of 4.5 × 10^-5^/bar was applied for pressure control. Covalent bonds are constrained to their equilibrium length by the LINCS algorithm (Hess et al., 1997). The integration steps of all simulations were set to 2 fs.

### PMF Calculations

For PC2 system, protein truncation was done on the equilibrated PIP_2_ and PC systems. Equilibrations were performed with unbiased MD simulations for 100 ns. Position restraints with a force constant of 1000 kJ/mol/nm^2^ were used to prevent protein translation during equilibration.

For PC2, the reaction coordinates were defined as the distance between the COM of Ser505 and Ser591 of PC2 and the whole head group of PC, PS (N*, PO* and GL* beads), PIP, PIP_2_ and PIP_3_ (RP*, PO* and GL* beads). 38 umbrella sampling windows were evenly spaced on the reaction coordinates between −0.7 and 3 nm with a force constant of 500 kJ/mol/nm^2^. Position restraints (400 kJ/mol/nm^2^ in the *xy* plane) were applied to the backbone beads of Ser505 and Ser591 in each subunit to prevent rotation and translation of the protein. The reaction coordinates are approximately parallel to the X axis of the simulation box and a weak positional restraint of 100 kJ/mol/nm^2^ was applied to the corresponding pulling groups of the lipids to limit its movements along the Y axis.

The PLUMED2 package (Tribello et al., 2014) was used to define the reaction coordinates and apply biasing to pull the lipids. WHAM was used to unbias all umbrella sampling simulations. For systems of all PIPs, the dissociation study was done on one of the four bound lipid molecules. For PC system, the protein was placed in pure PC bilayers, and one PC molecule situated in one of the binding sites was chosen to be pulled away from the proteins. For the PS system, the pulled PC in the PC system was replaced with a PS molecule, and a similar pulling protocol was applied. Simulations were run for 3 µs per window for all PIP systems and for 2 µs for the PC and PS systems. To calculate the energies, the first 800 ns of each window was discarded for all PIP systems, and the first 500 ns for the PC and PS systems.

### Simulation Visualisation and Analysis

Protein structures were visualised with VMD (Humphrey et al., 1996) or PyMOL (DeLano, 2002). Simulation trajectories were processed using GROMACS. Averaged lipid density maps were generated using the VMD Volmap plugin tool with three-dimensional grids every 0.1 nm.

### Protein Expression and Purification

The human PKD2 gene which encodes the PC2 (polycystin-2, PC2 or TRPP1) protein was purchased from the Mammalian Gene Collection (MGC, 138466; IMAGE, 8327731, BC112263). A truncated stable construct (hPC2: Pro185 - Asp723) was used, with a C-terminal purification tag containing a TEV cleavage site, a His_10_ sequence, and a FLAG tag, was cloned into the expression vector pFB-CT10HF-LIC (available from The Addgene Nonprofit Plasmid Repository). DH10Bac competent cells were used for the production of baculovirus. Recombinant baculoviruses were used to infect *Spodoptera frugiperda* (Sf9) insect cells, grown in 250 mL suspension in Sf900II serum free, protein-free insect cell medium with L-glutamine (Thermo Fisher Scientific) at 27°C, when cell density reached ∼2 × 10^6^/ml for virus amplification at 27 °C in 1 L shaker flasks. 1 L of Sf9 insect cells in Insect-XPRESS Protein-free Insect Cell Medium with L-glutamine (Lonza) in a 3 L flask was infected with 5 ml of the harvested P2 (second passage) viruses for 65 h at 27 °C. Cells were harvested 65 h post-transduction by centrifugation for 15 min at 1500 g and 4 °C.

Extraction buffer containing 50 mM HEPES, pH 7.5, 150 mM NaCl, 20 mM CaCl_2_, 5% glycerol and Roche protease-inhibitor cocktail was used to re-suspend the cell culture pellets to a volume of 50ml/L. Cells were lysed on ice with VCX 750 sonicator and 13mm probe (PRO Scientific Inc.) for 5 min, 3 sec on, 12 sec off and 35% amplitude. 1% (w/v) DDM was added to the cell lysis and incubated for 1 h at 4 °C. Cell debris was removed by centrifugation for 1h at 35,000*g* and 4 °C. To purify by immobilised metal affinity chromatography, the detergent-solubilised protein were batch bound to Co^2+^-charged Talon resin (Clontech) by gentle rotation at 4 °C for 1 h. The resin was washed with 15 column volumes of extraction buffer supplemented with 0.01% DDM and 30 mM imidazole, pH 8.0 and, to exchange the detergent, with another 15 column volumes of the same buffer replacing DDM with 0.035% UDM. hPKD2:185-723 protein was eluted from Talon resin with extraction buffer supplemented with 0.035% UDM and 400 mM imidazole. The eluted protein was divided into two batches. One was further purified via size-exclusion chromatography (SEC) with a Superose 6 increase 10/300GL column (GE Healthcare) pre-equilibrated with SEC buffer (0.035% UDM, 20 mM HEPES, pH 7.5, 200 mM NaCl and 20 mM CaCl_2_) for PIP strip experiments. The other was buffer exchanged into SEC buffer using a PD-10 column (GE Healthcare The protein was then treated with 0.4 units of bovine kidney α-L-fucosidase (Sigma) at 18 °C, overnight. The pH was adjusted to 7 before further enzymatic treatment at a ratio of 0.75:0.5:1 (w/w/w) TEV protease, PNGase F, PC2 for another 24 h at 18 °C. Reverse His-tag purification was performed to clear out the His_6_-tagged TEV protease and uncleaved PC2. The protein was concentrated to 0.5 ml with a 100-kDa-cutoff concentrator (Vivaspin 20, Sartorius), and further purified by SEC as above.

### PIP Strip Assay

PIP strip membranes (Thermo Fisher Scientific P23751) were blocked in 3% (w/v) fatty acid-free BSA (Sigma-Aldrich) in TBST (50 mM Tris-HCl, pH 7.5, 150 mM NaCl and 0.1% (v/v) Tween 20] for 1 h. The membranes were then incubated in the same solution with 0.5 μg/ml of detergent solubilised His_10_/FLAG-tagged PC2 overnight at 4 °C with gentle agitation. The membranes were washed 3 times over 30 min in fatty acid-free BSA-TBST. One more 3h-incubation with 0.5 μg/ml PC2 and subsequent wash steps were performed at 4°C. The membranes were incubated for 1h with 1:2000 dilution of HRP conjugated anti-FLAG monoclonal antibody (Thermo Fisher Scientific MA1-91878-HRP) at room temperature. Finally, the membranes were washed 6 times over 1 h in TBST, and the protein that was bound to the membrane by virtue of its interaction with phospholipid was detected by enhanced chemiluminescence.

### Cryo-EM Grid Preparation and Data Acquisition

18:0-20:4 PI(4,5)P_2_ and 18:1 PI(3,5)P_2_ lipid extract (Avanti) dissolved in chloroform was dried under an argon stream. 18:0-20:4 PI(4,5)P_2_ (∼ 0.5 mg/ml) and 18:1 PI(3,5)P_2_ (∼ 1 mg/ml) stock was prepared by resolubilising dried lipids in a buffer containing 20 mM HEPES (pH 7.5), and 150 mM NaCl via bath sonication for ∼1h. Purified PC2 was incubated with 18:0-20:4 PI(4,5)P_2_ at a molar ratio of 1:20 (PC2 tetramer: 18:0-20:4 PI(4,5)P_2_) and with 18:1 PI(3,5)P_2 2_ at a molar ratio of 1:40 (PC2 tetramer: 18:1 PI(3,5)P_2_) overnight at 4°C. The sample was cleared by centrifugation at 21,000g for 30 min. For cryo-EM, 3 μl of PC2 with PI(4,5)P_2_ sample at a protein concentration of ∼4 mg/ml or PC2 with PI(3,5)P_2_ sample at a protein concentration of ∼3.5 mg/ml was applied to glow-discharged Quantifoil 1.2/1.3 holey carbon 300 mesh copper grids. Grids were plunge frozen in liquid ethane using a Vitrobot Mark IV (FEI, Thermo Fisher Scientific) set to 5°C, 100% relative humidity, 3.5 s blotting time and −15 blotting force. Data for PC2 with PI(4,5)P_2_ were collected on a Titan Krios 300-kV transmission electron microscope equipped with a post-column Gatan image filter (GIF; 20eV slit width) operating in zero-loss mode and a Gatan K2 Summit direct electron detector camera at Central Oxford Structural Molecular Imaging Centre (COSMIC). Movies were captured for 0.4 s per frame over 8 s. The calibrated pixel size and dose rate were 0.822□Å/pix and ∼6.55 electrons/Å^2^/s, respectively (total dose 52.4e^-^/Å^2^). Images were collected in a defocus range between −1.0 and −3.0□µm under focus in 0.25 μm steps. Data for PC2 with PI(3,5)P_2_ were collected on a Titan Krios 300-kV transmission electron microscope at the Electron Bio-Imaging Centre (eBic, Diamond Light Source) equipped with a Gatan K2 Summit direct electron detector camera mounted behind a GIF and operated in zero-loss mode (0-20eV). Movies were captured for 0.2 s per frame over 7 s. The calibrated pixel size and dose rate were 0.816□Å/pix and ∼6.0 electrons/Å^2^/s, respectively (total dose 42e^-^/Å^2^). Images were collected in a defocus range between − 1.0 and −3.1□µm under focus in 0.3 μm steps.

### Image Processing

The beam-induced motion in the movies was corrected and frames were dose-weighted using MotionCor2 (Zheng et al., 2017). Aligned frames in each movie were averaged to produce a micrograph for further processing. The contrast transfer function (CTF) parameters were estimated using CTFFIND-4.1 (Rohou and Grigorieff, 2015). Micrographs with ice contamination or poor CTF cross correlation scores were discarded and the remaining micrographs were processed using RELION 3 (Kimanius et al., 2016). Particle picking was performed using Gautomatch (URL: http://www.mrc-lmb.cam.ac.uk/kzhang/).

For the dataset of PC2 with PI(4,5)P_2_, a set of 77,399 particles were picked from 1,597 micrographs and sorted into 2D classes. Representative 2D classes were used as templates for autopicking after low-pass filtering to 30 Å. A total of 147,001 particles were automatically picked. Three rounds of iterative 2D classification were performed in RELION to remove bad particles. A low-resolution reference model was generated *ab initio*. The initial model was used for the first round of 3D classification without symmetry imposed. A subset of 73,883 particles was used for subsequent 3D classification with C4 symmetry imposed. The data was then used for the first round of 3D ‘gold-standard’ refinement, which resulted in and initial reconstruction with a nominal unmasked resolution of 3.5 Å. Subsequent 3D classification was performed without further image alignments followed by iterative CTF refinement and Bayesian polishing. The final subset of particles was subjected to further auto-refinement in RELION, which converged at an unmasked resolution of 3.12 Å. RELION post-processing using unfiltered half maps and a soft-edged mask to exclude the region occupied by the detergent micelle yielded a final B-factor sharpened map (−84.56Å^2^) with a nominal resolution of 2.96 Å (FSC=0.143) (see SI Fig. S12 & S13).

For the dataset of PC2 with PI(3,5)P_2_, a set of 323,538 particles were initially picked from 3,353 micrographs and classified into 2D classes. Representative well resolved class averages were used as templates for reference-based particle picking after low-pass filtering to 30 Å. A total of 224,402 particles were automatically picked using GAUTOMATCH. Three rounds of iterative 2D classification were performed in RELION to remove bad particles. A low-resolution reference model was generated *ab initio*. The initial model was used for the first round of 3D classification without symmetry imposed. One further round of 3D classification with C1 symmetry was performed on a set of 97,892 particles from classes with the best estimated resolutions. A subset of 91,844 particles was used for subsequent 3D classification with C4 symmetry imposed. The data was then used for the first round of 3D ‘gold-standard’ refinement, which resulted in and initial reconstruction at a resolution of 4.1 Å. A final set of 37,297 particles was selected from 3D classification performed without further image alignments. Iterative CTF refinement and Bayesian polishing were performed prior to a final auto-refinement procedure in RELION, which converged at an unmasked resolution of 3.66 Å. Subsequent post-processing, using a soft-edged mask that excluded the detergent micelle, produced a 3.4 Å resolution map. Reported resolutions were based on a FSC threshold of 0.143. Local resolutions across the whole map were estimated using RESMAP (SI Figure S14) (Kucukelbir et al., 2014).

### Model Building

The previously published model of apo PC2 (PDB 5K47) was used as an initial model and fitted into the cryo-EM map of lipid-bound PC2. The model was manually adjusted in Coot (Emsley and Cowtan, 2004). In the original PDB 5K47 structure, the disulphide bond between Cys311 and Cys334 appeared to be reduced. In both PIP complexes, there is clear density showing that a disulphide bond is formed between these cysteines and has been built in both structures. One spherical density corresponding to the size of a hydrated sodium or calcium ion is clearly present below the selectivity filter in each of the PIP complexes maps with the density deeper in the central pore in PI(3,5)P_2_ map compared to PI(4,5)P_2_ map. No such density is present in the equivalent location in map EMD-8354 (corresponding to PDB id 5T4D; [Shen, 2016 #6850]). The difference in the ionic species between the 5T4D/EMD-8354 and our preparation is that our buffer contains calcium. Therefore, calcium ions were into the ion density in both of our structures. A series of *B*- factor sharpened maps were used to guide model building. Final models of both structures were globally refined and minimised in real space against the RELION3 C4 post-processed, automatically *B*-factor sharpened maps with NCS constraints, secondary structure and rotamer restraints imposed and no Ramachandran restraints applied using the *phenix.real_space_refine* module in PHENIX (Adams et al., 2010). The refinement protocol was validated by taking the final refined models, applying a random shift of up to 0.3Å to the atomic coordinates and then refining the resultant shifted model against the halfmap1. Model-to-map FSCs were calculated using *phenix.mtriage* against halfmap1 (FSCwork), halfmap2 (FSCfree) and the full map (FSCsum).

## Acknowledgements

This work was funded by grant to MSPS from BBSRC, EPSRC, Wellcome, and UCB. The SGC is a registered charity (number 1097737) that receives funds from AbbVie, Bayer Pharma AG, Boehringer Ingelheim, the Canada Foundation for Innovation, Genome Canada, Janssen, Lilly Canada, Merck KGaA, Merck & Co., Novartis, the Ontario Ministry of Economic Development and Innovation, Pfizer, São Paulo Research Foundation-FAPESP and Takeda, as well as the Innovative Medicines Initiative Joint Undertaking ULTRA-DD grant 115766 and the Wellcome Trust (106169/Z/14/Z). A.G. is supported by a BBSRC and Pfizer funded studentship.

We acknowledge the Oxford Particle Imaging Centre (OPIC) for providing access to electron microscopes for grid screening. We also acknowledge the Central Oxford Structural Microscopy and Imaging Centre (COSMIC) for providing access to electron microscopes for grid screening and data collection, and thank Chitra Shintre (SGC), Errin Johnson and Adam Costin (COSMIC) for assistance with grid screening and data collection. We acknowledge Diamond Light Source for access and support of the cryo-EM facilities at the UK’s national Electron Bio-imaging Centre (eBIC; under BAG proposal em14856), funded by the Wellcome Trust, MRC and BBRSC. We acknowledge the use of the UCSF Chimera package from the Resource for Biocomputing, Visualisation, and Informatics at the University of California, San Francisco (supported by NIGMS P41-GM103311).

We thank all members of the SGC Biotech team and the IMP1 group for assistance with this work. We thank Brian Marsden and David Damerell, James Bray, James Crowe and Chris Sluman for bioinformatics support. Our thanks also to Jan Domański for his help with PMF calculations.

