## Supplemental Figures and Table for "Lipid Interactions of a Ciliary Membrane TRP Channel: Simulation and Structural Studies of Polycystin-2 (PC2)"

**Supplementary Information**

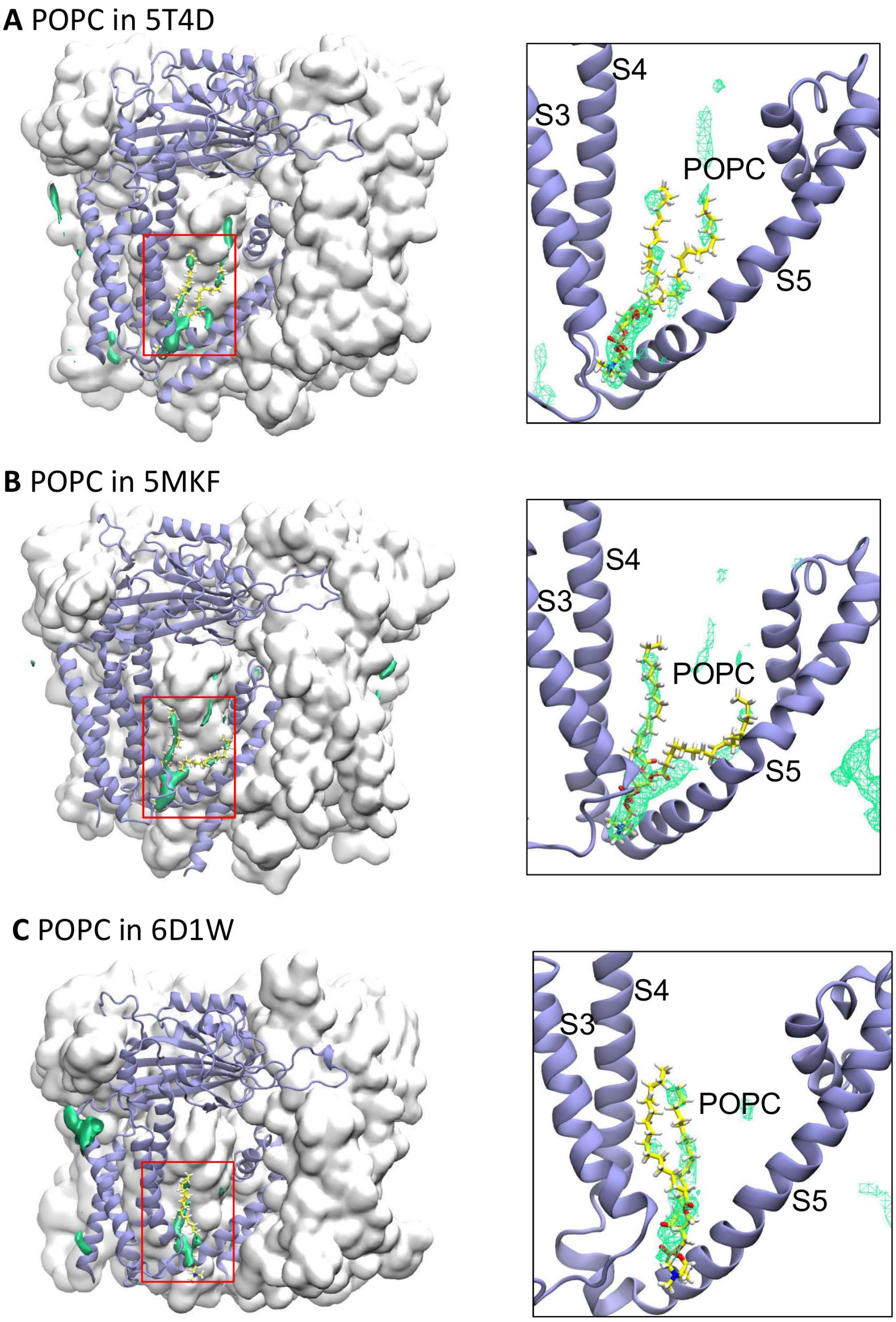


*Figure S1:*

Probability density of phosphatidylcholine (PC) in atomistic simulations based on three different PC2 cryo-EM structures: **A** 5T4D, **B** 5MKF, and **C** 6D1W. In each case in the left panel PC2 is shown as a grey surface, viewed perpendicular to the central pore axis, with one subunit depicted as a pale purple cartoon. Green isocontour surfaces represent a high probability of occurrence of phospholipid molecules. In the right panels, corresponding zoomed in views (red box) of the S3/S4/S5 pocket and the high phospholipid occurrence density are shown with a PC molecule taken from a simulation snapshot.

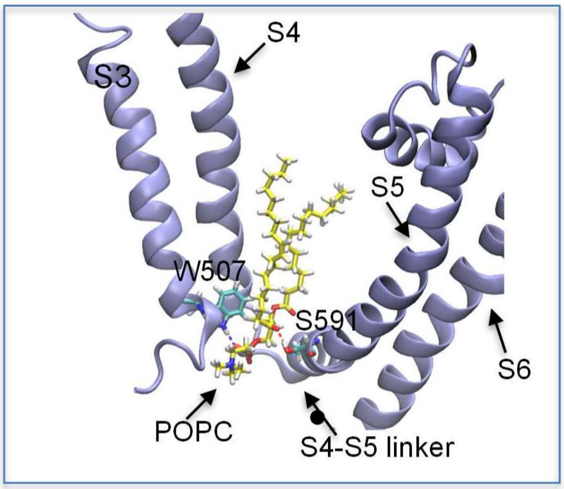


*Figure S2:*

PC interaction with PC2 at the possible lipid binding site identified in the atomistic simulations. S3-S6 of PC2 (taken from a simulation snapshot of 5K47) is shown as pale purple cartoon with a bound PC molecule shown as yellow sticks. Residues Trp507 and Ser591 which formed the most hydrogen bonds between the lipid and protein during the simulations are shown as blue sticks.

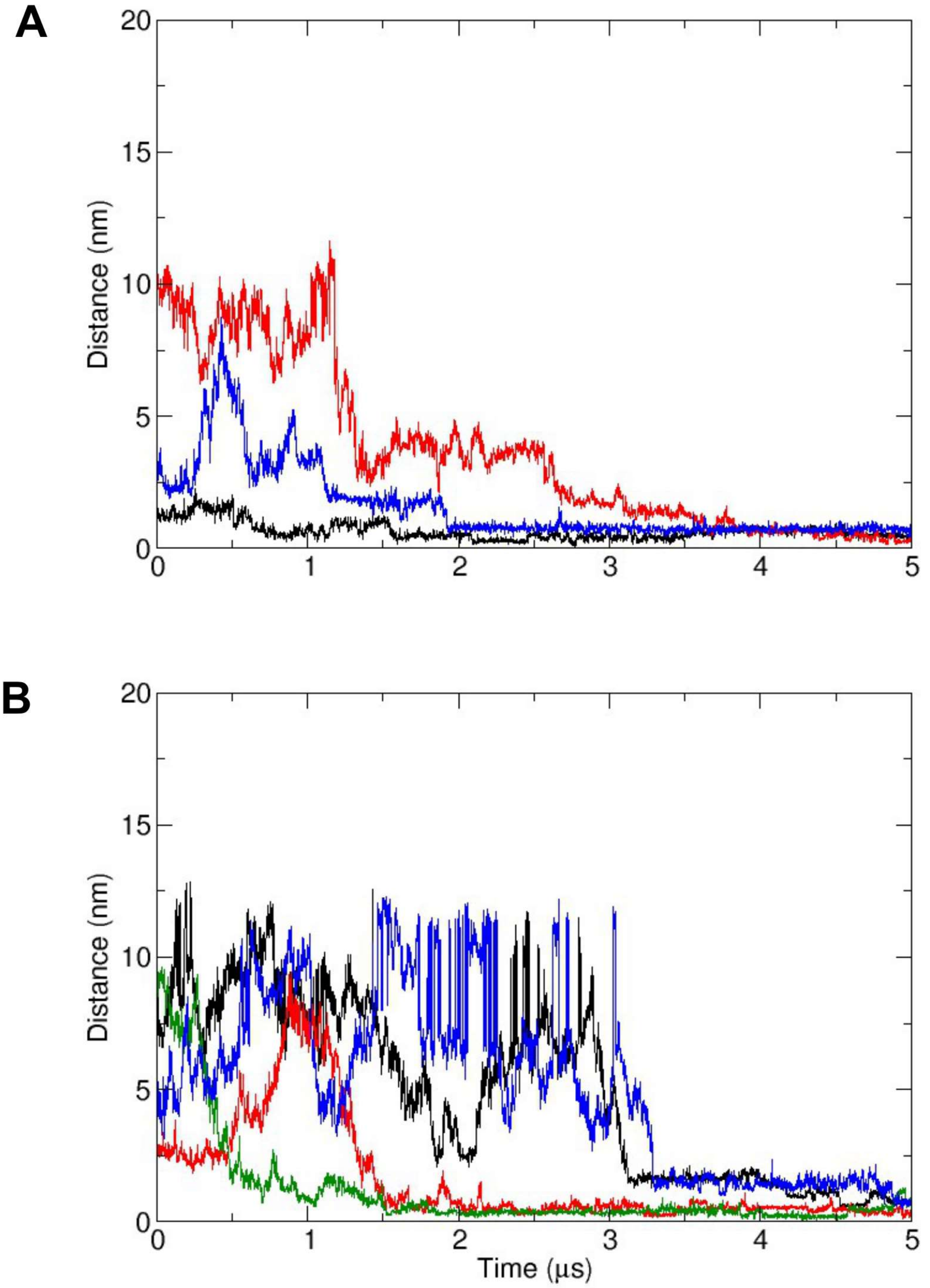


*Figure S3:*

$PIP_2$ binding in two repeats of a coarse-grained (CG) simulations of PC2 (structure 5K47) in an *in vivo* mimetic mixed lipid bilayer (also see Fig. 2A of the main paper for the third simulation of this triplicate). The distances from their binding site on PC2 of $PIP_2$ molecules are shown as functions of time. Distances are measured from the centre of mass of the headgroup of each $PIP_2$ molecule to the centre of mass of residues S505 and S591 of the site to which that lipid molecule eventually binds. The different coloured lines correspond to the different $PIP_2$ molecules.

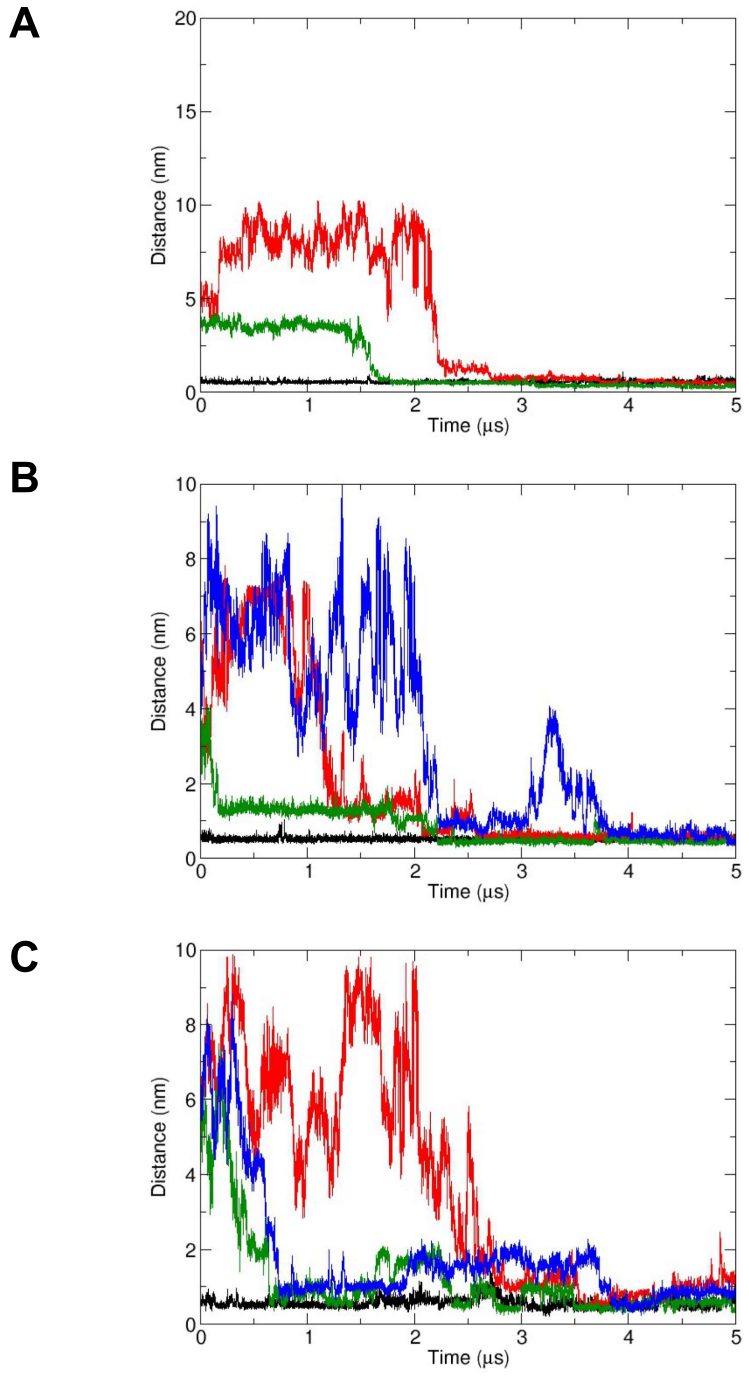


*Figure S4:*

$PIP_2$ binding in three repeats of CG simulations of a gain-of-function Phe604Pro mutant of PC2 (6D1W) in an *in vivo* mimetic mixed lipid bilayer. Other details are as for Figure S3.

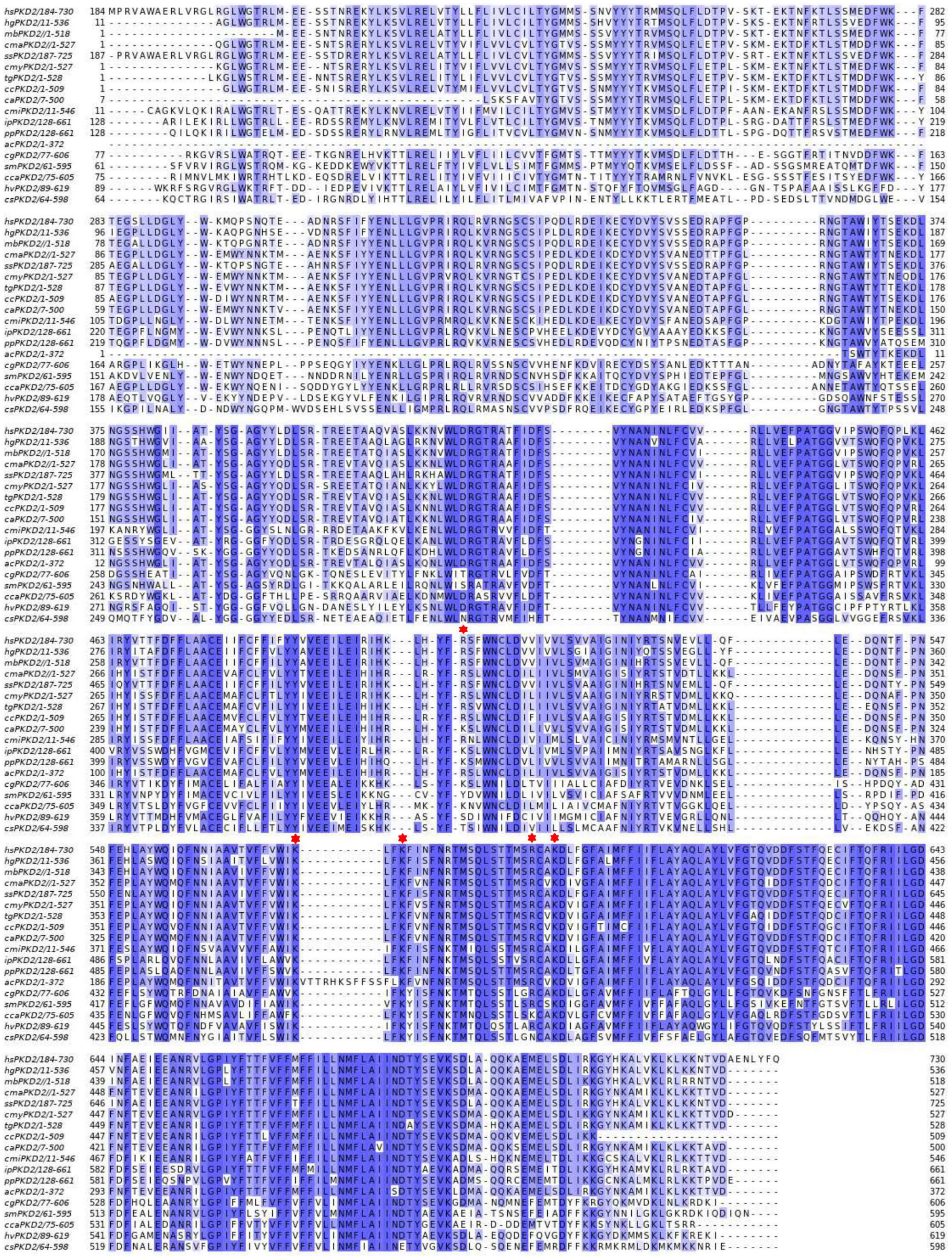

*Figure S5:*

Sequence alignment for PC2 from 18 different species. The sequence of human PC2 (M184-Q730) (hsPKD2) is aligned with the corresponding sequences from *H. glaber* (hg), *M. brandtii* (mb), *C. macqueenii* (cma), *S. scrofa* (ss), *C. mydas* (cmy), *T. guttatus* (tg), *C. canorus* (cc), *C. anna* (ca), *C. milii* (cmi), *I. punctatus* (ip), *P. prolifica* (pp), *A. chloris* (ac), *C. gigas* (cg), *S. mimosarum* (sm), *C. candelabrum* (cca), *H. vulgaris* (hv), *C. sinensis* (cs). Residues Arg504, Lys572, Lys575, Arg592 and Lys595, identified to coordinate the $PIP_2$ headgroup in human PC2, are highlighted by red stars.

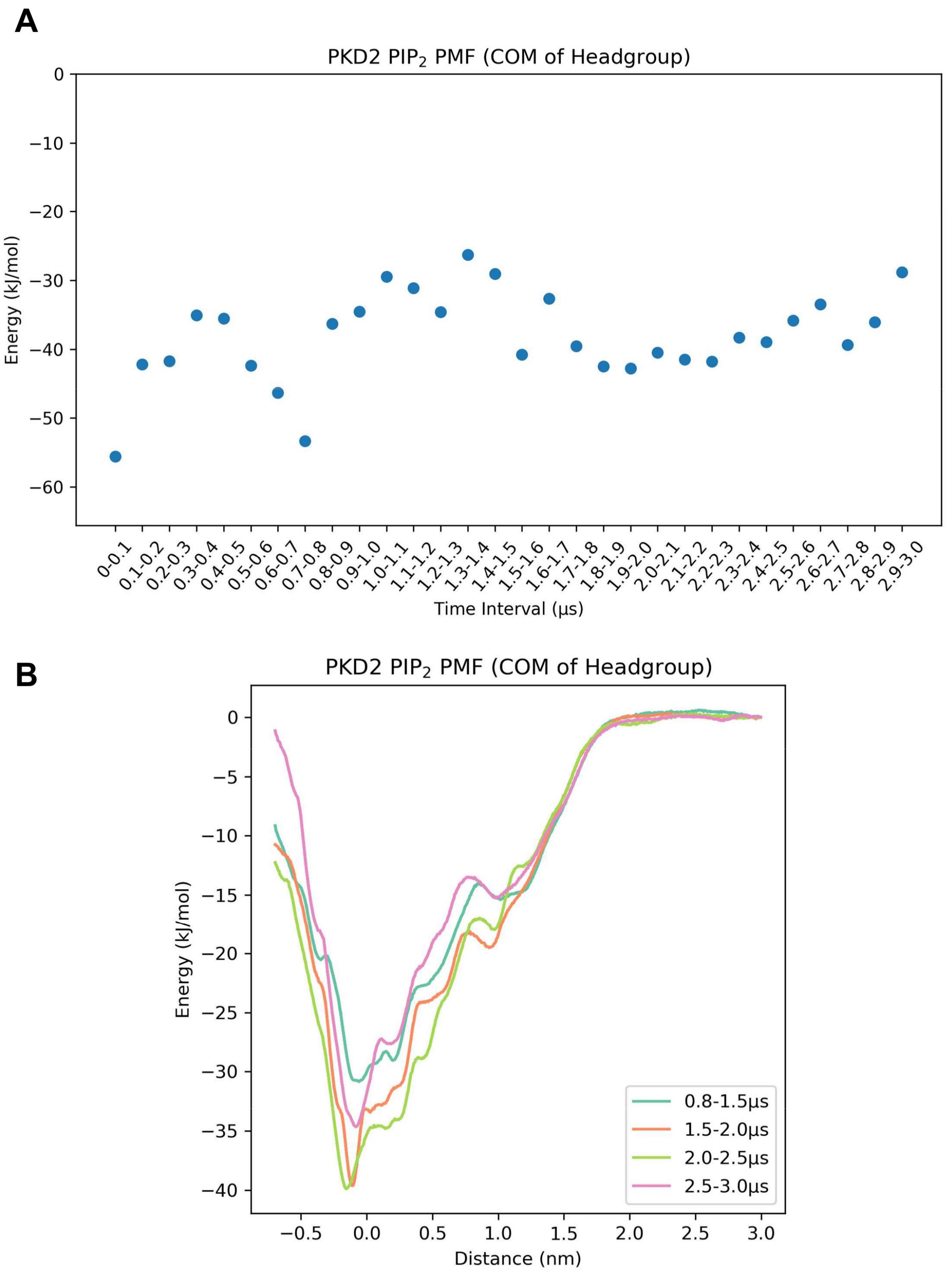


*Figure S6:*

Convergence of PMF calculations of $PIP_2$ binding to PC2. **A** Convergence of a PMF calculation for $PIP_2$ binding was assessed by calculating the binding free energy (measured as the energy well depth) derived from non-overlapping 100 ns segments of the simulations. **B** PMF profiles calculated with non-overlapping 500 to 700 ns time segments are shown. The first 800 ns of each window were excluded from the PMF calculation.

A

B

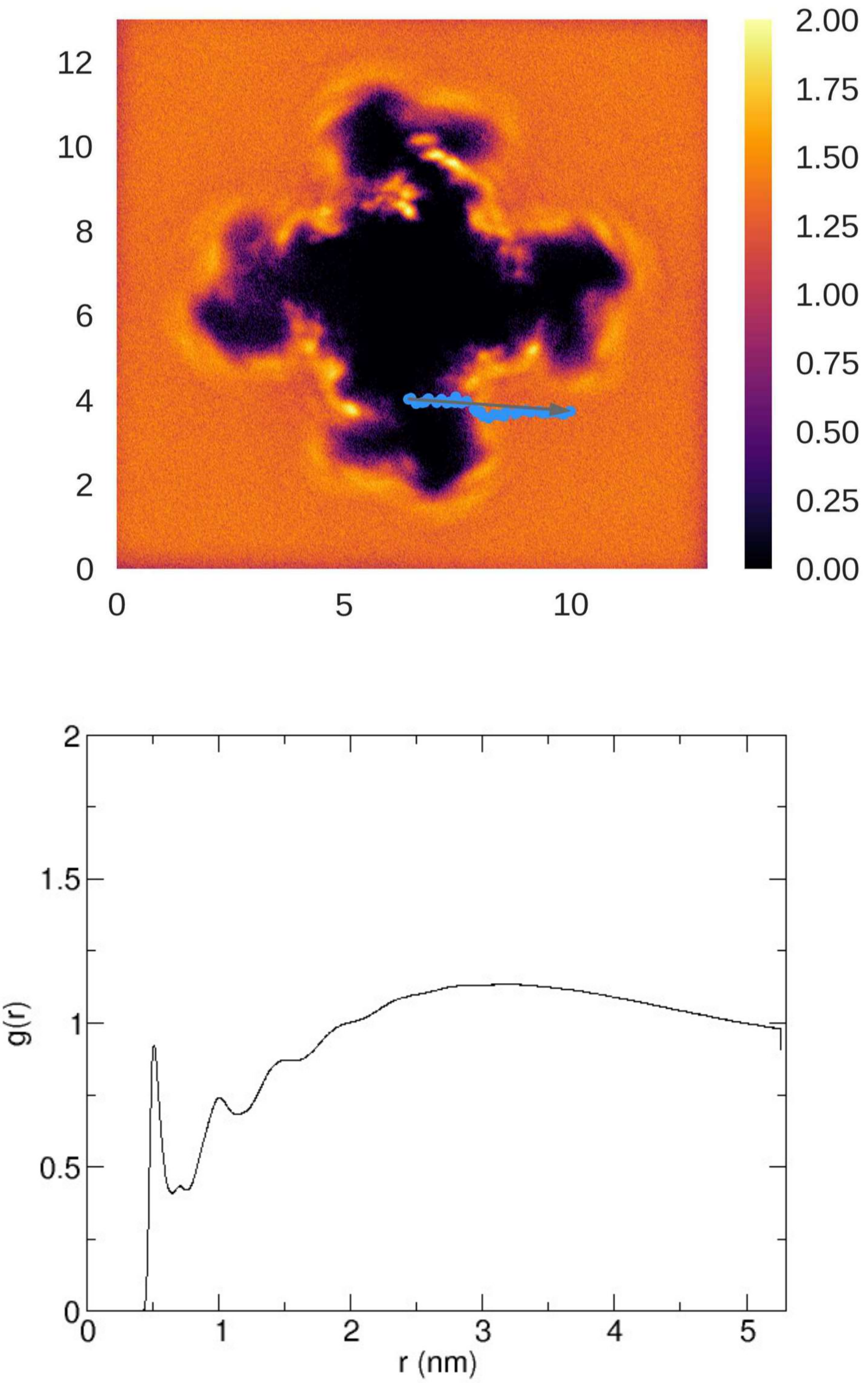


*Figure S7:*

PC distribution in the cytoplasmic leaflet of the bilayer around PC2 in a $PIP_2$ PMF simulation. **A** PC density around PC2 is plotted with the $PIP_2$ reaction (i.e. pulling) coordinates indicated as blue dots. The pulling direction is shown with ochre arrow. **B** The radial distribution of PC around PC2.

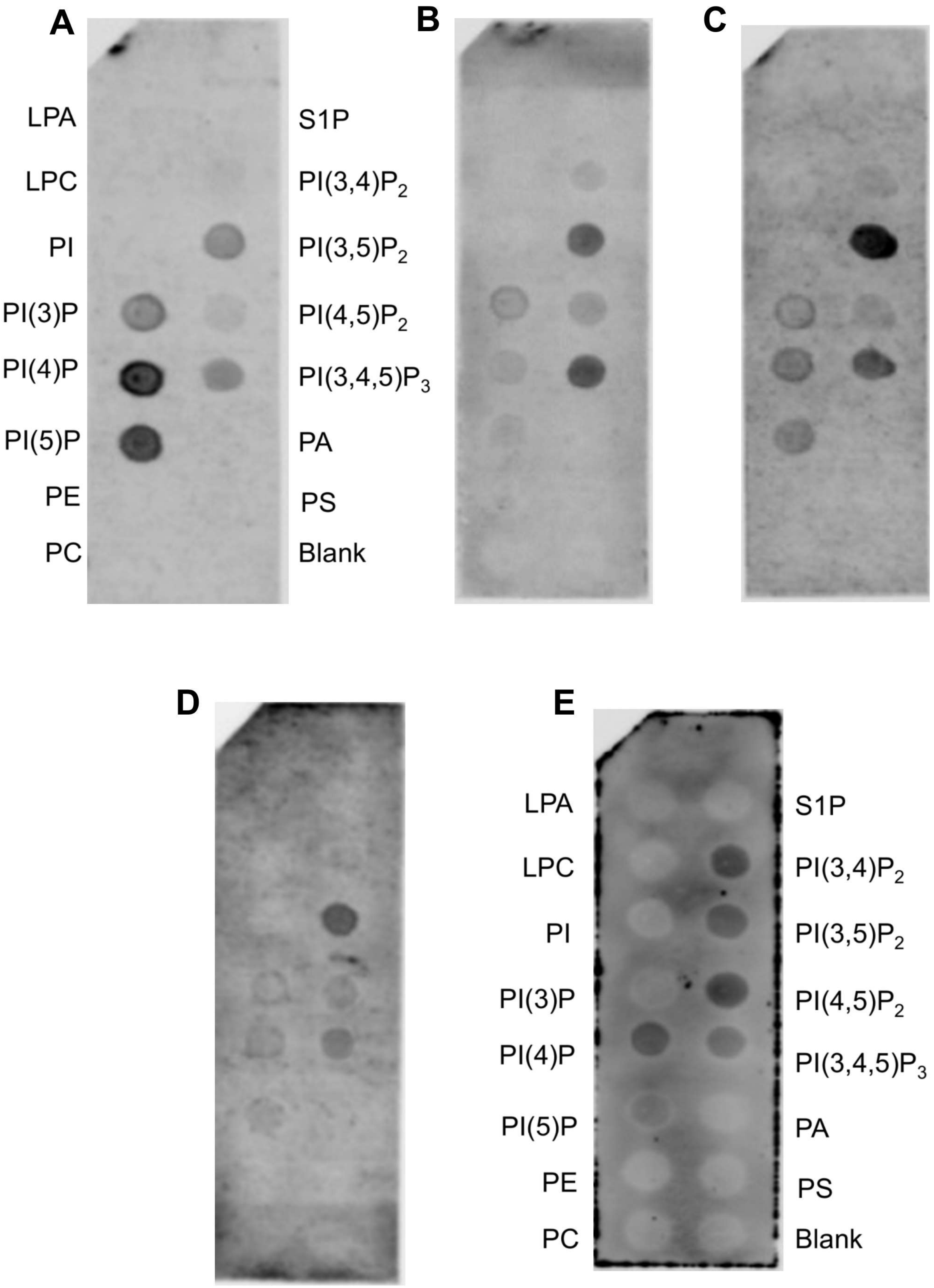


*Figure S8:*

Biological repeats of PC binding to PIP strips. PC2 expressed in different batches of either GNT-HEK (**A**) or Sf9 (**B - E**) cells were used to repeat the PIP strip experiments. Although binding intensities vary between the different repeats, the bound lipid species are broadly consistent.

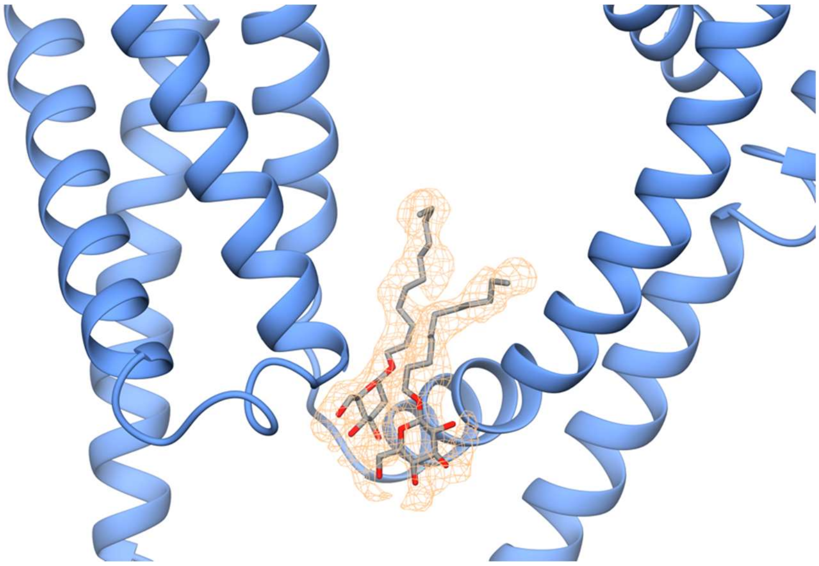

*Figure S9:*
Zoomed-in view of the lipid binding site density in the 3.0 Å structure with PI(4,5)P2 present. Two UDM molecules are modelled between S3, S4 and S5 helices. Detergent/lipid density is shown in orange.

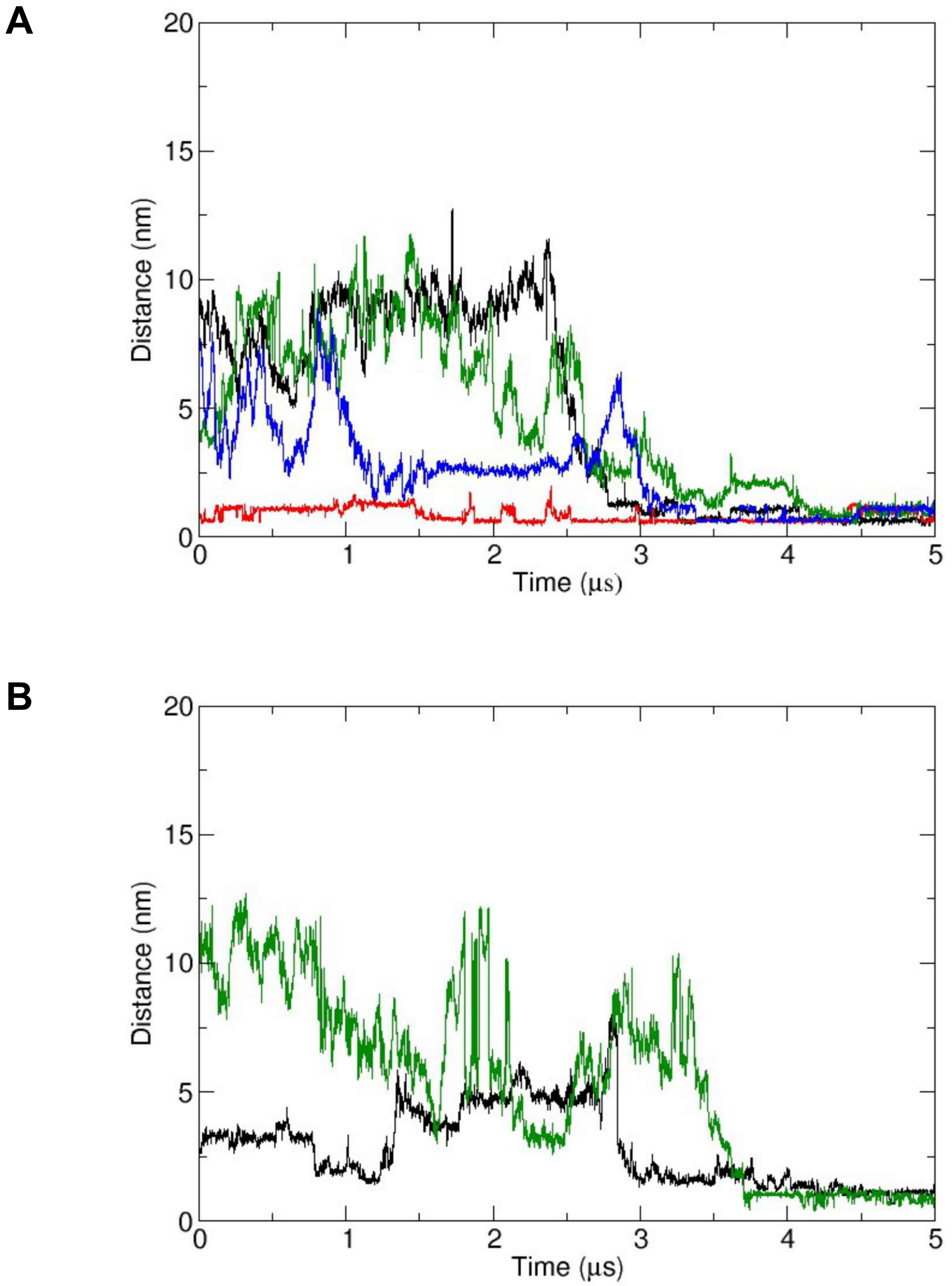


<u>*Figure S10:*</u>
Cholesterol binding in two repeats of CG simulations of PC2 (structure 5K47) in an *in vivo* mimetic mixed lipid bilayer. Distances from the binding site as function of time for cholesterol molecules are shown. The distance shown is from the centre of mass of each cholesterol molecule to the centre of mass of Gln557 and Asn560 at the site to which that lipid molecule eventually binds. Different coloured lines correspond to the different cholesterol molecules.

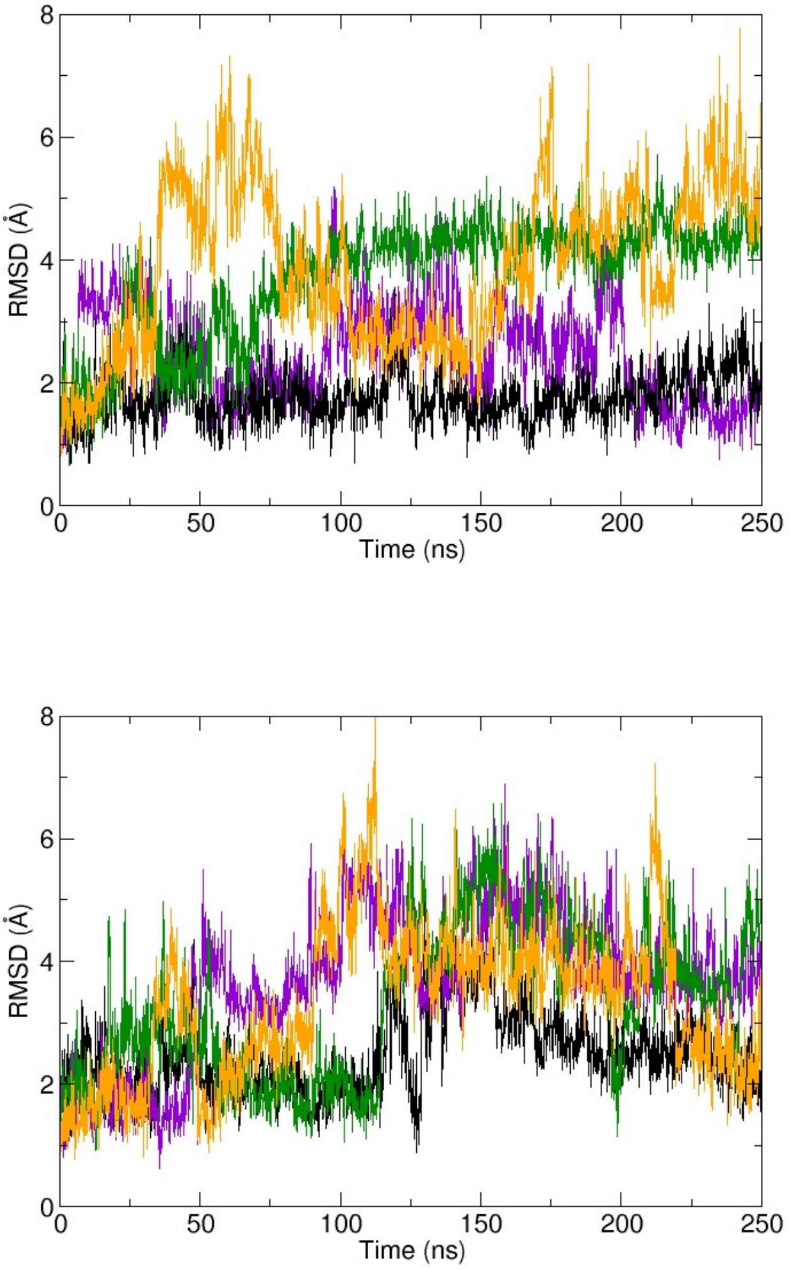


*Figure S11:*

Two repeats of atomistic simulations of cholesterol-bound PC2, with the cholesterol molecule initially built into the cryoEM density. RMSD vs. time for the four cholesterol molecules bound to PC2 during 250 ns atomistic MD simulations are shown. Different coloured lines correspond to the different cholesterol molecules.

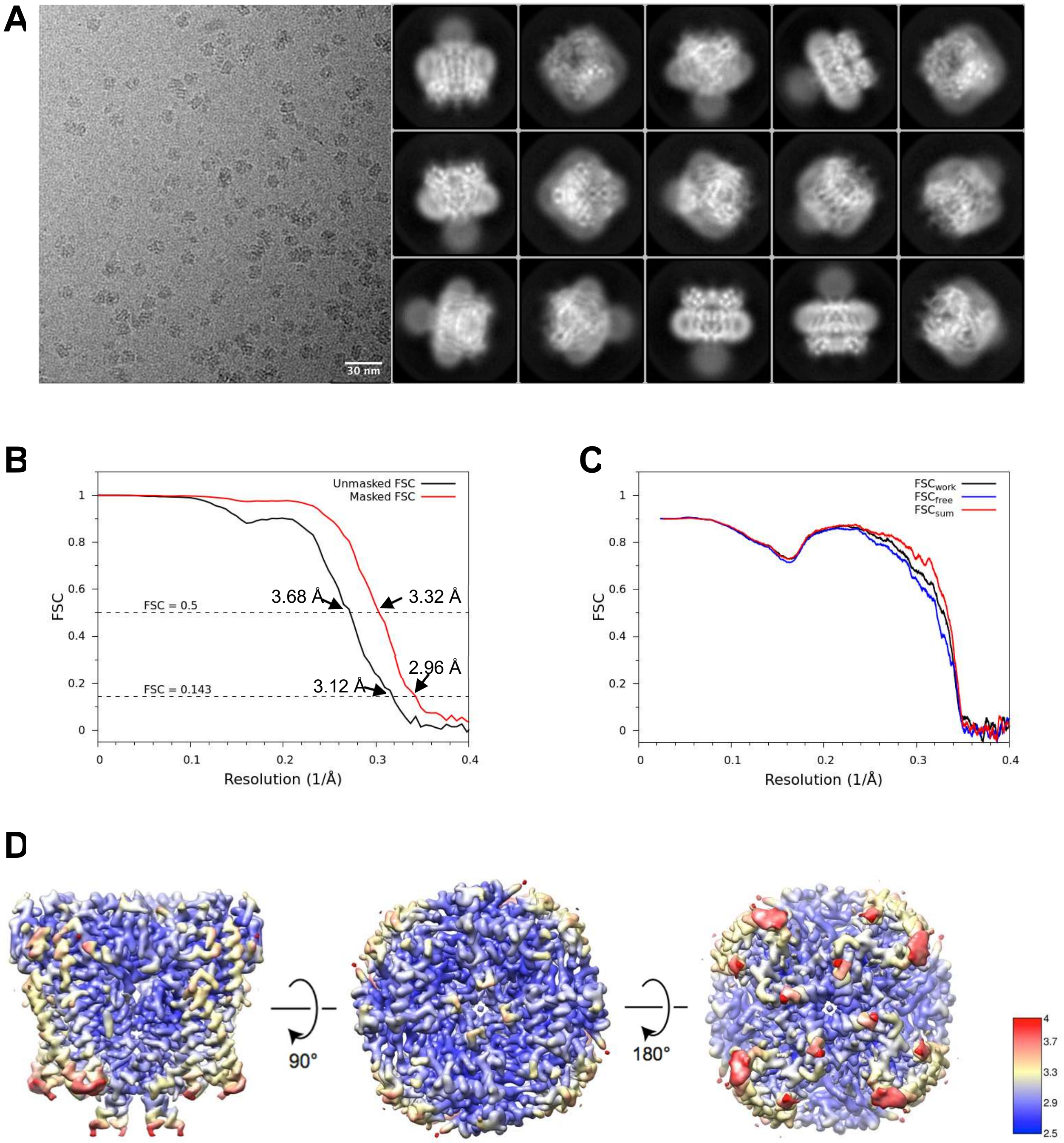


*Figure S12:*

PC2 structure with PI(4,5)$P_2$ in detergent. (**A)** A representative raw micrograph and 2D classes of detergent-solubilized PC2 with PI(4,5)$P_2$. (**B**) Fourier shell coefficient (FSC) curves of the masked (red) and unmasked (black) maps. FSC of 0.143 and 0.5 are indicated with dotted lines. (**C**) FSC curves for cross validation comparing the model to half maps 1 (black) and 2 (blue), and to summed map (red dotted). (**D**) Local resolution of the final density map in three different orientations on a scale of 2.5 Å (blue) to 4 Å (red).

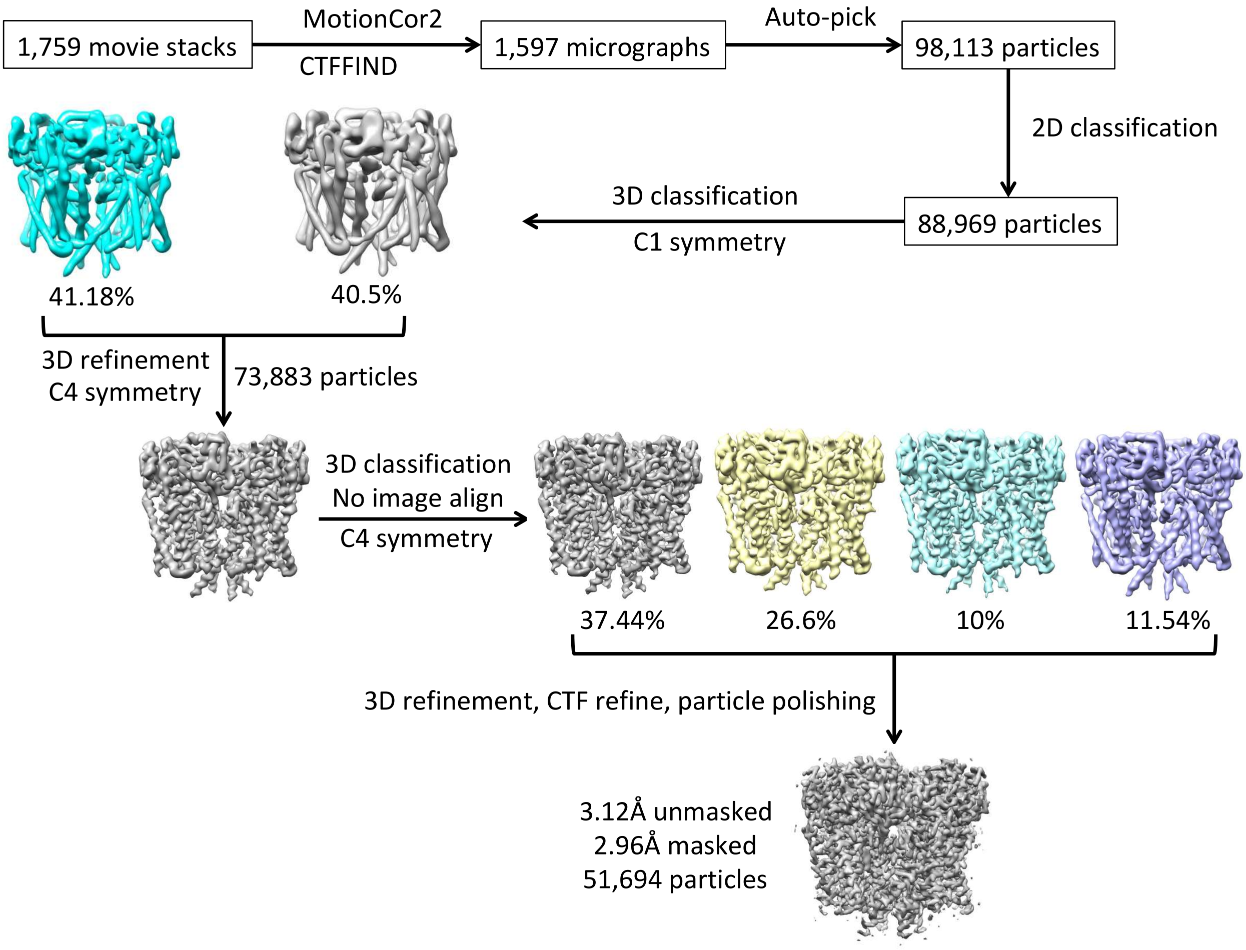


*Figure S13:*

Data processing for the PC2 dataset with PI(4,5)$P_2$ in detergent. Workflow of processing the images of PC2 with PI(4,5)$P_2$ to 3 Å resolution.

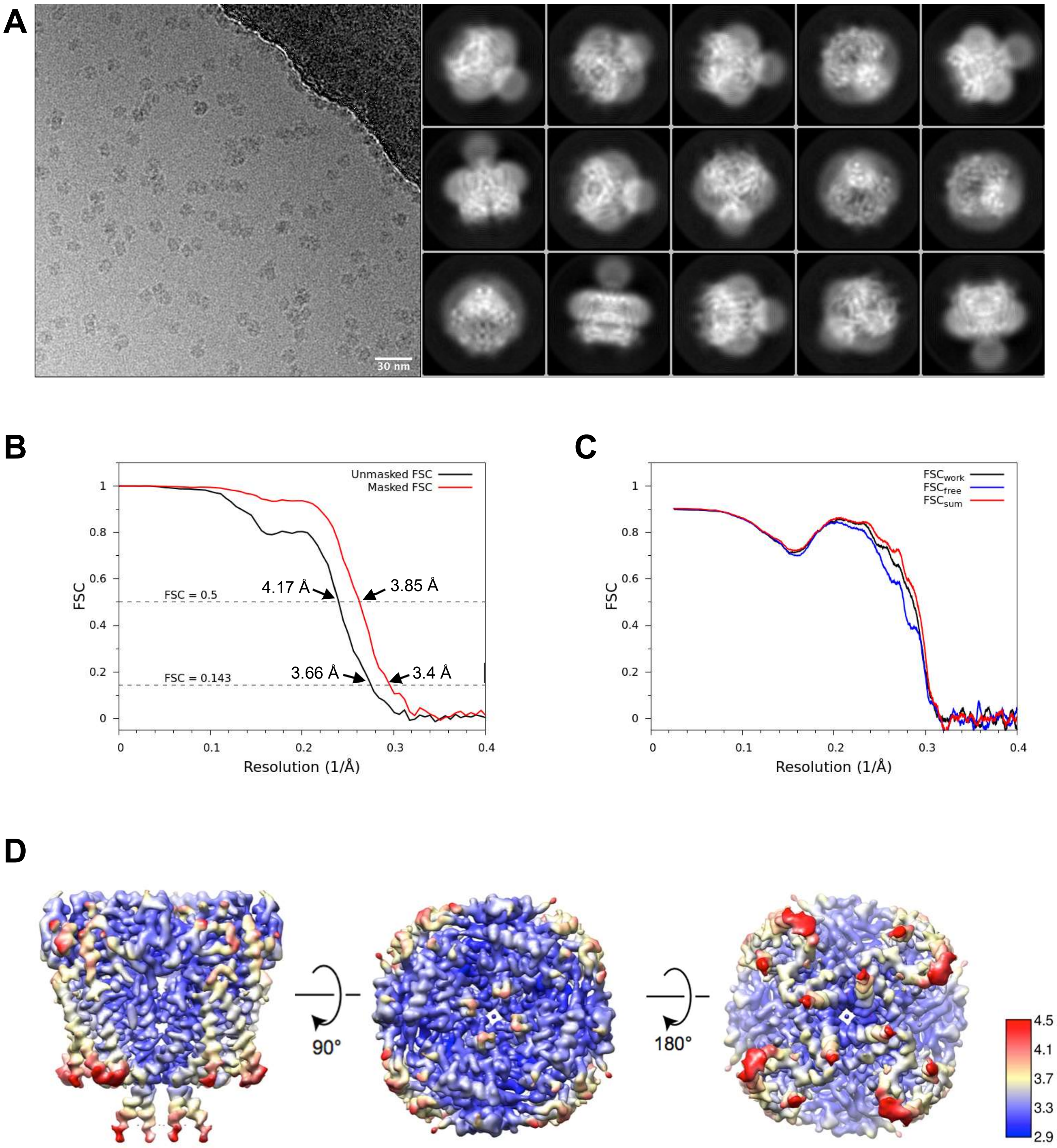


*Figure S14:*

PC2 structure with PI(3,5)$P_2$ in detergent. (**A**) A representative raw micrograph and 2D classes of detergent-solubilized PC2 with PI(3,5)$P_2$. (**B**) Fourier shell coefficient (FSC) curves of the masked (red) and unmasked (black) maps. FSC of 0.143 and 0.5 are indicated with dotted lines. (**C**) FSC curves for cross validation comparing the model to half maps 1 (black) and 2 (blue), and to summed map (red dotted). (**D**) Local resolution of the final density map in three different orientations on a scale of 2.9 Å (blue) to 4.5 Å (red).

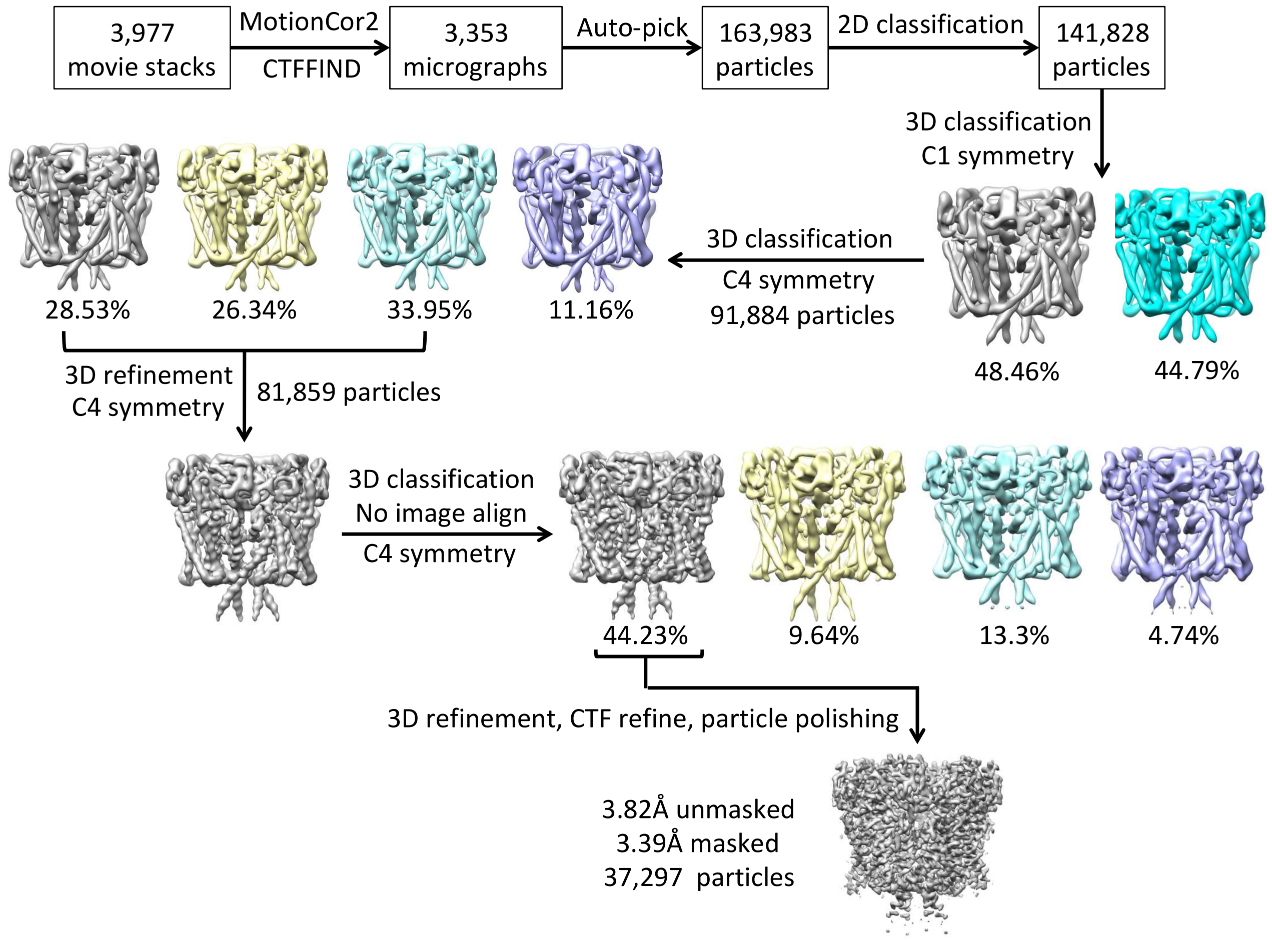


*Figure S15:*
Data processing for the PC2 dataset with PI(3,5)$P_2$ in detergent. Workflow of processing the images of PC2 with PI(3,5)$P_2$ to 3.4 Å resolution.

**Extended Data Table 1 |** Cryo-EM data, refinement and model statistics.

| **Cryo-EM data** | PI(4,5)$P_2$ | PI(3,5)$P_2$ |
|---|---|---|
| **Data Collection:** | | |
| Voltage (kV) | 300 | 300 |
| Defocus range (μm) | -3 to -1 in 0.25 increments | -3.1 to -1 in 0.3 increments |
| Pixel size (Å) | 0.822 | 0.816 |
| Electron dose ($e^-$/Å$^2$) | 52.4 | 42.05 |
| Dose rate ($e^-$/Å$^2$/s) | 6.55 | 6.0 |
| Number of micrographs (used) | 1,597 | 3,353 |
| Particles (initial)[1] | 98,113 | 163,983 |
| Particles (final, %) | 51,694 (52.7) | 37,297 (22.7) |
| **Reconstruction:** | | |
| Symmetry | C4 | C4 |
| Resolution (unmasked, Å)[2] | 3.12 | 3.66 |
| Resolution (masked, Å)[2] | **2.96** | **3.39** |
| Map sharpening *B*-factor (Å$^2$) | -84.56 | -108.53 |
| **Model composition:** | | |
| Protein atoms | 15,668 | 15,612 |
| Other | 1,249 | 869 |
| **Refinement:** | | |
| Resolution (Å) | **3** | **3.4** |
| Map sharpening factor (Å$^2$) | -84.56 | -108.53 |
| Fourier Shell Correlation (FSC)[3] | 0.8485 | 0.8309 |
| **Rms deviations:** | | |
| Bonds (Å) | 0.011 | 0.009 |
| Angles (°) | 1.002 | 0.875 |
| **Molprobity validation:** | | |
| Clashscore, all atoms | **3.72** | **3.63** |
| Molprobity score | 1.43 | 1.61 |
| Ramachandran Plot (% favoured) | 96.01 | 92.66 |
| Ramachandran Plot (% allowed) | 3.57 | 6.71 |
| Ramachandran Plot (% outliers) | 0.42 | 0.63 |

[1]Particles after one cycle of 2D classification to remove non-particles
[2]Based on FSC 0.143 threshold
[3] CC_mask from *phenix_real_space_refine*
